# TRAF6 initiates inflammatory signaling via organizing membraneless cytoplasmic condensates

**DOI:** 10.1101/2023.06.19.545655

**Authors:** Jia Wang, Xincheng Zhong, Hang Yin

## Abstract

The tumor necrosis factor receptor-associated factor 6 (TRAF6) is a central molecule in multiple signaling pathways, i.e. the TNF receptor, the Toll-like receptors (TLRs), and interleukin-1 receptor (IL-1R) pathways. Upon pathogen associated molecular patterns (PAMPs) or damage associated molecular patterns (DAMPs) stimulations, TRAF6 activates downstream of NF-κB signaling. However, the precise mechanism of how TRAF6 activates downstream molecules remains unclear.

Here, we demonstrate that TRAF6 acts as a sensor for upstream signals to initiate liquid-liquid phase separation (LLPS) of itself and downstream proteins, forming membraneless condensates. Subsequent recruitment, enrichment and activation of downstream effectors in the condensates lead to robust inflammatory signal transduction. The multivalent interactions mediated by its RING domain, zinc finger domain 1, and coiled-coil domain mediates the LLPS process. Forced phase separation of TRAF6 induced NF-κB activation. Disruption of TRAF6 phase separation abrogates activation of NF-κB signaling. Overall, we uncover the spatial organization of molecules by TRAF6 through phase separation as a subcellular platform to activate inflammatory signaling. Targeting TRAF6 phase separation hold promises for therapeutics aiming autoimmune diseases, inflammation and cancers.

## Introduction

TRAF6 is firstly identified as a mediator of IL-1R-mediated activation of NF-κB (Cao *et al*., 1996). TRAF6 acts downstream of multiple receptor families, such as TLRs (O’Neill *et al*, 2013), tumor growth factor-β receptors (TGFβR) (Hamidi *et al*, 2017), T-cell receptors (TCR) (Sun *et al*, 2004), and The tumor necrosis factor (TNF) receptor superfamily member 11a (TNFRSF11a, also known as RANK) (Darnay *et al*, 1999). At the physiological level, TRAF6 plays essential roles in innate and adaptive immunity (Wu & Arron, 2003), embryogenesis (Naito *et al*, 2002), bone metabolism (Armstrong *et al*, 2002) and various cancer developments (Starczynowski *et al*, 2011; Rezaeian *et al*, 2017; Li *et al*, 2020).

As an unconventional E3 ubiquitin ligase (Deng *et al*, 2000), TRAF6 catalyzes the synthesis of K63-linked polyubiquitin chains and transfers ubiquitin to its substrates. Upon stimulations, TRAF6 forms complexes with Ubc13/UBE2N and Uev1A/UBE2V1 to generate K63-linked polyubiquitin chains, which are then attached to target proteins, such as TAK1, inhibitory kappa B kinase γ(IKKγ, or NEMO), and TRAF6 itself (Deng *et al*, 2000; Wang *et al*, 2001; Kanayama *et al*, 2004; Ea *et al*, 2006; Wu *et al*, 2006). The binding of TAB2 or TAB3 with polyubiquitin chains promotes the activation of TAK1 through self-phosphorylation (Ea *et al*, 2006; Wu *et al*, 2006). Next, TAK1 phosphorylates inhibitory kappa B kinase β (IKKβ), inducing I kappa B kinase alpha(IκBα) phosphorylation. IκBα then will undergo K48 polyubiquitination coupled degradation, resulting in translocation of NF-κB complex (Liu & Chen, 2011). TAK1 can also phosphorylate mitogen-activated protein kinase kinase (MAPKK), causing the activation of c-Jun N-terminal kinase (JNK) and p38 pathways(Chen *et al*, 1995; Brown *et al*, 1995; Wang *et al*, 2001).

LLPS is a process by which macromolecules such as proteins or nucleic acids condense into liquid droplet-like structures. LLPS drives the formation of many membraneless compartments, such as nucleoli, Cajal bodies, stress granules, postsynaptic densities, and signaling puncta (Banani *et al*, 2017). Several recent reports have shown LLPS involves in the regulation of immunological signal transduction. For example, phase separation of T cell receptors in Jurkat T cells (Su *et al*, 2016); DNA-induced liquid phase condensation of cGAS (Du & Chen, 2018); and during B cell activation, SLP65, CIN85, and lipid vesicles phase separated into droplets (Wong *et al*, 2020). However, whether the NF-κB signaling pathway is regulated by LLPS has not been investigated.

In this study, we show that TRAF6 undergoes LLPS both at the cellular level and in vitro. Interestingly, membraneless condensates initiated by TRAF6 involve components of ubiquitin chain synthesis for efficient production of long ubiquitin chains. TAK1, TAB2 and NEMO are also concentrated into condensates to accelerate their phosphorylation and ubiquitination processes, finally resulting in strong activation of inflammatory signaling. In summary, we reveal a higher level organization of TRAF6 and its downstream interacting proteins during PAMPs and DAMPs recognition, which efficiently promotes signal transduction and amplification.

## Results

### TRAF6 forms droplet-like structures in cells under stimulations

TRAF6 acts as a molecular hub connecting upstream and downstream molecules in TLR, TCR, RANK, IL17R, and TGFβRI signaling (Walsh *et al*, 2015). Multivalent interactions are the crucial driving force behind phase separation. We hypothesized that TRAF6 might took advantage of LLPS for efficient signal transduction. Thus EGFP-tagged TRAF6 either at the N- or the C-terminal were overexpressed in HeLa cells. Both N- and C-terminal EGFP-tagged TRAF6 formed droplet structures in the cytosol with the N-terminal EGFP-tagged TRAF6 forming larger droplets of several microns (Figure 1A). Then we used N-terminal EGFP tagged TRAF6 in the subsequent experiments. Next, the fluidity of these droplets was assessed by fluorescence recovery after photobleaching (FRAP). The results showed that the fluorescence of TRAF6 droplets recovered ranging from 30% to 80% within 20 minutes (Figure 1B and C), indicating that they form gel-like condensates with relatively low fluidity. Additionally, two droplets adjacent to each other could fuse together into a larger one (Figure 1D). The above results revealed the common phase separation characteristics of TRAF6.

**Figure 1.**
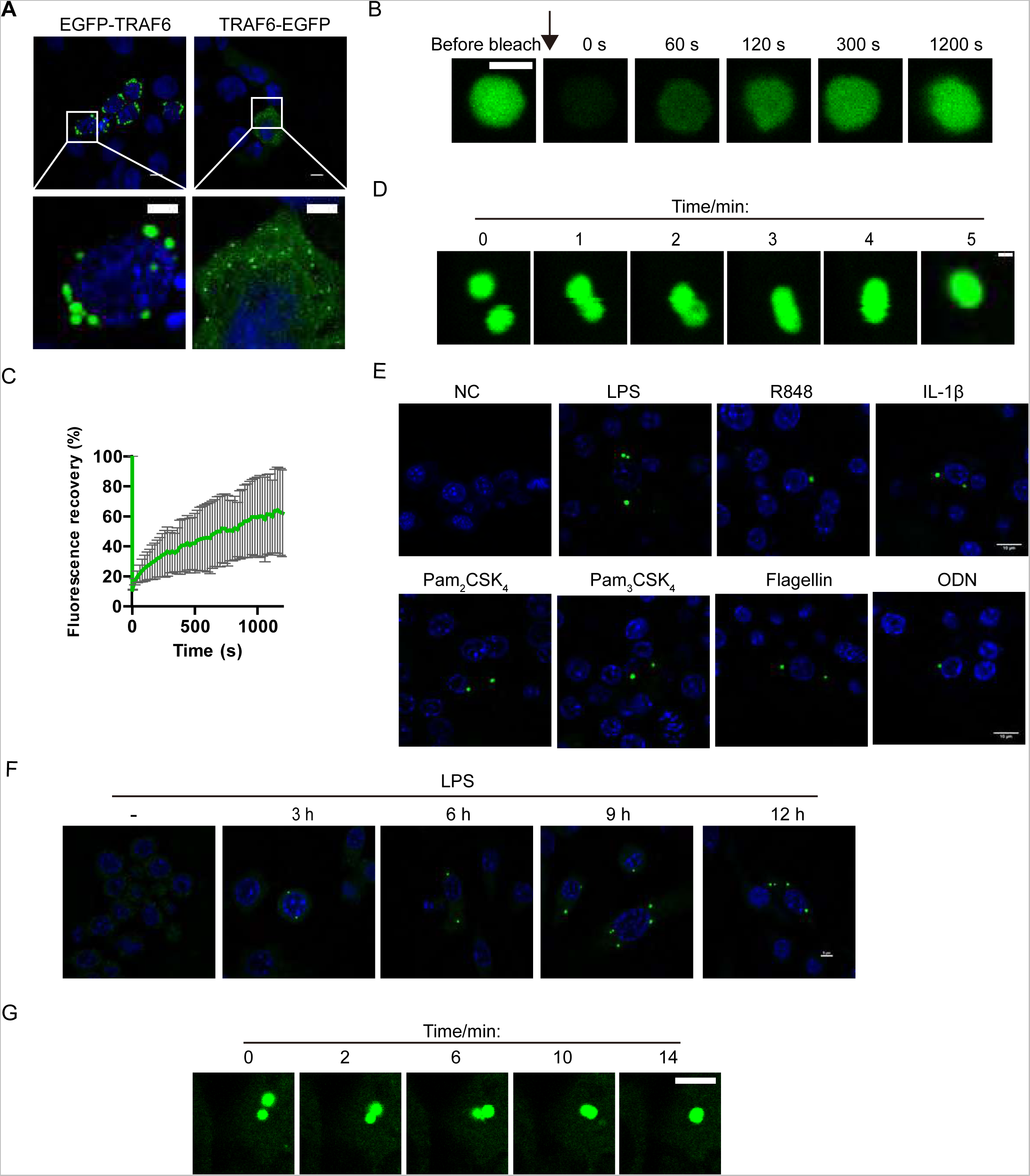
TRAF6 condensate into droplets in cells. A. EGFP-TRAF6 and TRAF6-EGFP formed condensates in HeLa cells.
B. FRAP of EGFP-TRAF6 and time lapse tracking of fluorescence recovery.
C. Statistics analysis of EGFP-TRAF6 condensates’ FRAP.
D. Fusion of EGFP-TRAF6 condensates.
E. Stimulation of Raw264.7^EGFP-TRAF6^ cells with LPS (100 ng/ml), R848 (2 μg/ml), Pam_2_CSK_4_ (100 ng/ml), Pam_3_CSK_4_ (100 ng/ml), Flagellin (100 ng/ml) and ODN (2 μg/ml) for 12 hours, then the cells were fixed and observed under NIKON AIR confocal microscope.
F. Raw264.7^EGFP-TRAF6^ cells were stimulated with LPS (100 ng/ml) for 3, 6, 9 and 12 hours separately, then fixed and observed under confocal microscope.
G. Raw264.7^EGFP-TRAF6^ cells were stimulated with LPS (100 ng/ml) for 1 hours, then cultured in a live cell chamber, recorded in a time lapse manner. Data information: Scale bars, 10 μm in (A, E and F) and 2μm in (B, D and G).

To further investigate whether TRAF6 can phase separate under physiological conditions, we constructed a Raw264.7 cell line (Raw264.7^EGFP-TRAF6^) stably expressing low levels of EGFP-TRAF6 (Figure EV1A). In the resting state, EGFP-TRAF6 exhibited a diffused cellular distribution in Raw264.7^EGFP-TRAF6^ cells (Figure 1E). However, upon stimulation with agonists of TLRs or IL-1R, i.e., LPS (TLR4), R848 (TLR7/8), Pam_2_CSK_4_ (TLR2/6), Pam_3_CSK_4_ (TLR1/2), Flagellin (TLR5), ODN (TLR9) and IL-1β (IL1R), EGFP-TRAF6 condensed into droplets in the cytosol (Figure 1E). The sizes of droplets were in the range of micrometers. These results indicate that TRAF6 droplets formation was regulated by upstream signals under physiological conditions. Observation at different time points demonstrated that TRAF6 droplets could be stable for several hours (Figure 1F). In addition, when Raw264.7^EGFP-TRAF6^ cells were stimulated with LPS, TRAF6 droplets also could fuse with one another (Figure 1G). Collectively, these data demonstrate that TRAF6 can phase separate into condensates at the cellular level.

### TRAF6 undergoes phase separation in vitro

TRAF6 is composed of four domains: the RING domain, zinc finger (ZF) domain 1-4, the coiled-coil domain, and the TRAF-C domain. From the perspective of structure, the RING domain interacts with Ubc13/UB2N, and the zinc finger domain (ZF) 1 interacts with ubiquitin (Ub) (Yin *et al*, 2009). The coiled-coil domain is responsible for TRAF6 oligomerization and promotes TRAF6 ubiquitin ligase activity (Hu *et al*, 2017). The TRAF-C domain forms mushroom-like trimers to interact with upstream proteins such as IRAK1 and RNAK (Ye *et al*, 2002). From our data, it can be concluded that TRAF6 assembles into higher order complexes rather than just oligomerization.

To further dissect which domains are required for TRAF6 droplets formation, a series of EGFP-tagged TRAF6 truncation and deletion plasmids based on published structures and ALFA FOLD 2 predictions were constructed and overexpressed in HeLa cells (Figure EV2A). The images exhibited that neither the N-terminal 49 amino acids nor the TRAF-C domain were required for TRAF6 condensate formation (Figure EV2A). Additionally, none of the individual domain (the RING domain, ZF1 - ZF4 domain, the coiled-coil domain, and the TRAF-C domain) alone could form condensates in cells (Figure EV2A). Interestingly, overexpression of TRAF6(50-350) proteins in HeLa cells produced many droplets comparable to that by full-length TRAF6 (Figure EV2A). These data suggest that a combination of the RING domain, ZF1 - ZF4 and the coiled-coil domain is sufficient for TRAF6 condensation.

To determine which domain is essential for TRAF6 phase separation, deletion plasmids encompassing amino acids 50-349 were constructed and overexpressed in HeLa cells (Figure 2A). We found that deleting ZF2, ZF3, or ZF4 separately had no significant effect on droplet formation for both full-length TRAF6 and TRAF6(50-349) (Figure 2B). However, deleting either ZF1 or the coiled-coil domain strongly impaired droplet formation (Figure 2B). Deletion RING in full-length TRAF6 or TRAF6(50-349) led to protein aggregates in the nucleus. The above results suggest that both ZF1 and the coiled-coil domain are essential for inducing condensate formation. Previous study reports that the coiled-coil domain is responsible for TRAF6 oligomerization and constant interaction with Ubc13/UBE2N(Hu *et al*, 2017). Here, we demonstrate that the coiled-coil domain plays an important role in promoting phase separation.

**Figure 2.**
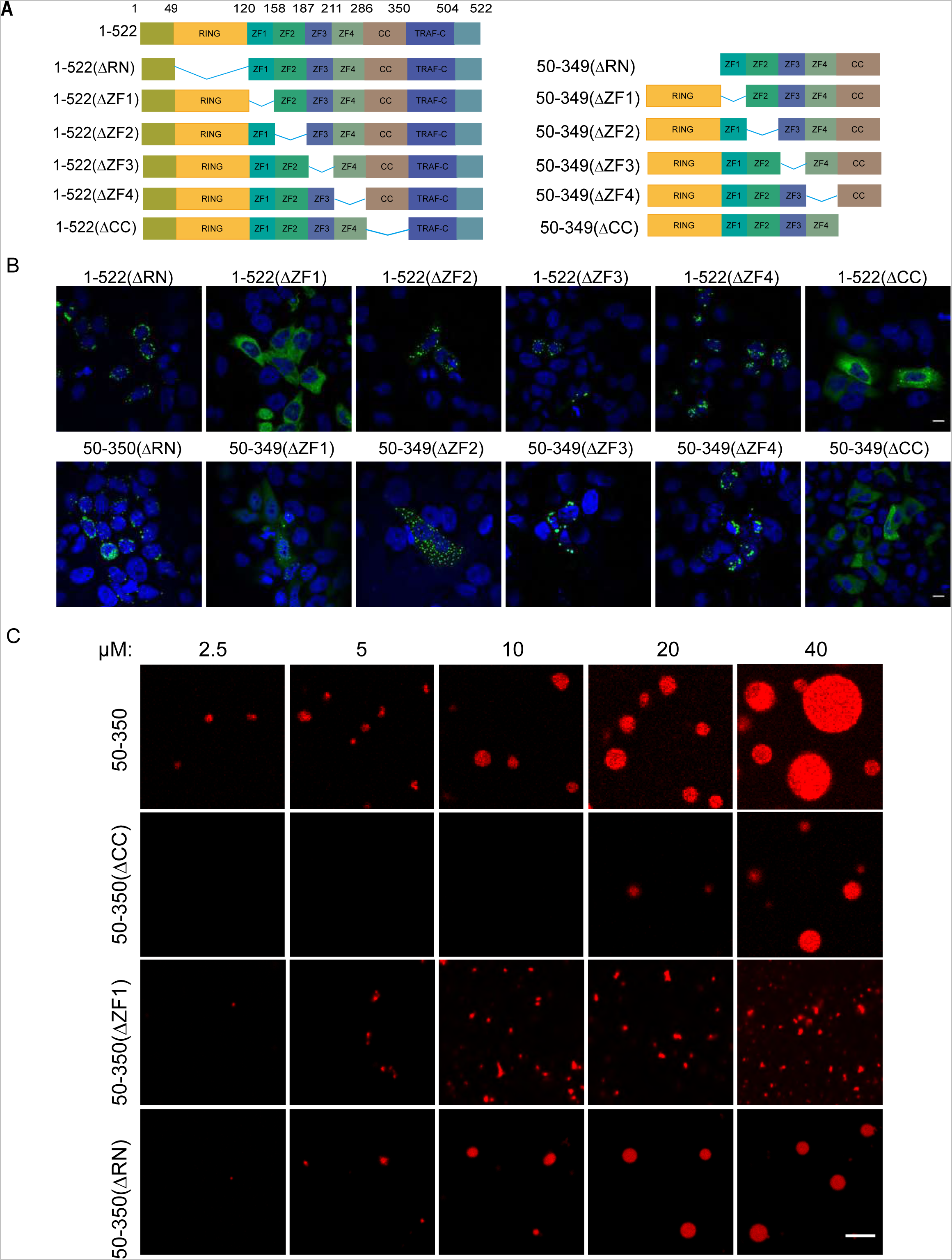
Multivalent interactions promote TRAF6 phase separation. A. Schematic diagram of TRAF6 deletions and truncations.
B. Representative images of TRAF6 deletions and truncations overexpressed in HeLa cells.
C. Representative in vitro reconstitution images of TRAF6(50-350), TRAF6(ΔCC), TRAF6(ΔZF1) and TRAF6(ΔRING) in a gradient manner. Data information: Scale bars, 10 μm in (B) and 5 μm in (C).

Next, to validate our cellular results, we reconstituted TRAF6 phase separation in vitro. Purified TRAF6(50-350), TRAF6(50-350ΔCC), TRAF6(50-350ΔZF1) and TRAF6(50-350ΔRN) proteins were mixed in a reconstitution buffer at different concentrations. Droplets appeared at 2.5 μM for TRAF6(50-350) proteins (Figure 2C). The sizes of liquid droplets increased with the increasing protein concentrations, indicating that TRAF6 phase separation happens in a concentration-dependent manner. For TRAF6(50-350ΔCC) proteins, smaller droplets only appeared above a concentration of 20 μM, indicating the importance of the coiled-coil domain in lowering the threshold concentration for phase separation. Compared to TRAF6(50-350), both TRAF6(50-350ΔZF1) and TRAF6(50-350ΔRN) proteins produced smaller condensates at the same concentrations (Figure 2C), suggesting that deleting either ZF1 or the RING domain weakens TRAF6’s ability to phase separate. The different abilities of TRAF6(50-350ΔRN) proteins to form condensates in cells and in vitro may be due to its strong interaction with endogenous TRAF6 through coiled-coil domains. FRAP data showed that TRAF6(50-350) formed condensates with low mobility (Figure EV2B and C).

Both in cells and in vitro results consistently demonstrate that TRAF6 can undergo LLPS and that the RING domain, ZF1, and the coiled-coil domain play fundamental roles.

### TRAF6 phase separation promotes K63 poly-ubiquitin chain synthesis

TRAF6 is a RING type E3 ubiquitin ligase that catalyzes the synthesis and transfer of K63 ubiquitin chains in synergy with the ubiquitin-activating enzyme (E1), the Ubc13-Uev1A (UBE2N/UBE2V1) ubiquitin-conjugating enzyme 2 (E2) complex, and Ub. To investigate whether TRAF6 condensates contain ubiquitin chains, we performed immunofluorescence staining for Ub and K63 Ub in HeLa cells overexpressing EGFP-TRAF6. Ub and K63 Ub signals showed obvious enrichment at TRAF6 condensates (Figure 3A), suggesting that LLPS promotes the synthesis of K63 Ub chains.

**Figure 3.**
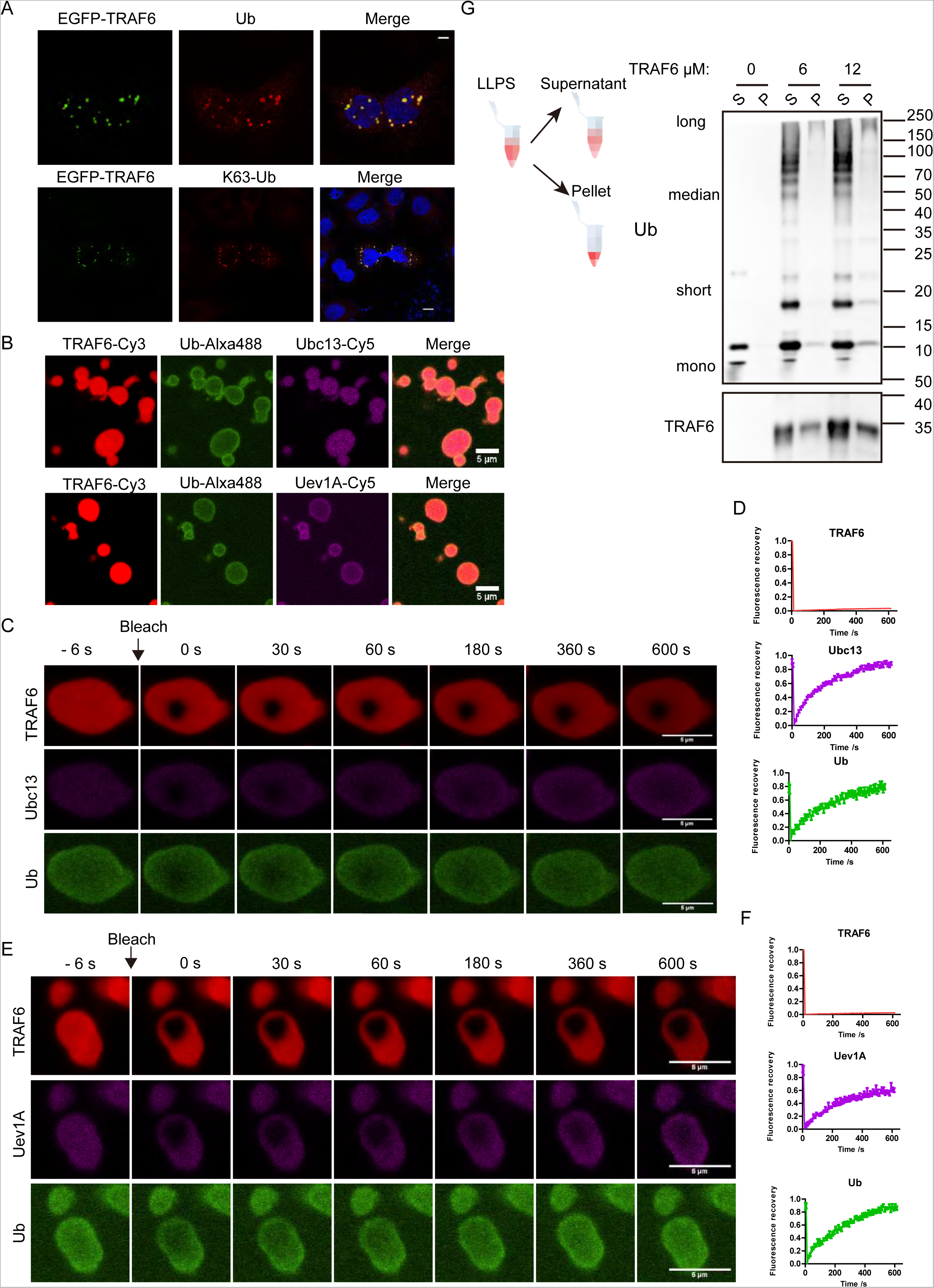
Components of ubiquitin chain synthesis were involved in TRAF6 LLPS. A. Upper panel, immunostaining of ubiquitin in EGFP-TRAF6 overexpressed HeLa cells; lower panel, immunostaining of K63 ubiquitin in EGFP-TRAF6 overexpressed HeLa cells.
B. Upper panel, in vitro LLPS of TRAF6, ubiquitin and Ubc13; lower panel, in vitro LLPS of TRAF6, ubiquitin and Uev1A.
C. Representative images of fluorescence recovery of a TRAF6, Ubc13 and Ub condensate.
D. FRAP analysis assaying the exchange kinetics of each protein between the TRAF6 condensates and the solution in (C).
E. Representative images of fluorescence recovery of a TRAF6, Uev1A and Ub condensate.
F. FRAP analysis assaying the exchange kinetics of each protein between the TRAF6 condensates and the solution in E. The concentrations of TRAF6, Ubc13, Uev1A, Ub and E1 were 20 μM, 1 μM, 1 μM, 50 μM and 0.1 μM separately in (B-F). ATP concentration was 2 mM in (B-F).
G. Sedimentation assays to analyze the retention of ubiquitin chains after LLPS and ubiquitin synthesis. The concentrations of Ubc13, Ube2V1, Ub, E1 and ATP were the same with that in B-F. Data information: Scale bars, 10 μm in (A), 5 μm in (B, C and E).

Next, to further confirm our hypothesis, we reconstituted K63 Ub synthesis in vitro using purified proteins, including TRAF6(50-350), E1, Ubc13/UBE2N, Uev1A/UBE2V1 and UB in a buffer containing ATP. Immunoblotting against Ub showed successful synthesis of polyubiquitin chains (Figure EV3A). Imaging of Alexa488-labeled Ub, Cy5-labeled Ubc13/UBE2N, and Cy3-labeled TRAF6 in the reconstitution system revealed that both Ub and Ubc13/UBE2N were recruited into TRAF6 droplets (Figure 3B). Similarly, Uev1A/UBE2V1 was also enriched in TRAF6 condensates (Figure 3B). Since none of UB, Ubc13/UBE2N, or Uev1A/UBE2V1 could undergo LLPS (data not shown), TRAF6 acted as a driver to initiate this phase separation process, while E1, UB, Ubc13/UBE2N and Uev1A/UBE2V1 acted as client proteins. In FRAP experiments, TRAF6 fluorescence hardly recovered 10 minutes after bleaching, indicating that it forms gel-like condensates. However, the fluorescent signals of Ubc13/UBE2N, Uev1A/UBE2V1, and UB almost completely restored within 10 minutes after bleaching (Figure 3C-F). These results suggest that TRAF6 forms low-fluidity condensates resembling scaffolds while ubiquitin chain synthesis components can move and interact freely into and out of condensates, thus accelerating the reaction.

We hypothesize that TRAF6 phase separation could form a micro-reactor to promote K63 Ub chain synthesis. In the sedimentation experiment, short and median (2-8) ubiquitin chains were mostly found in the supernatant, while long polyubiquitin chains were trapped in pellets (Figure 3G). Long unanchored polyubiquitin chains have been reported to be important for the phosphorylation and activation of TAK1 (Wang *et al*, 2001). TAB2 and TAB3 activate the NF-κB signaling through binding to polyubiquitin chains (Kanayama *et al*, 2004). Thus, LLPS of TRAF6 and subsequent polyubiquitin chain synthesis may provide a reaction platform to gather downstream proteins for efficient ubiquitination and phosphorylation.

### TRAF6 phase separation recruits and activates downstream proteins

It is known that after TRAF6 activation and polyubiquitin synthesis, a series of proteins are activated, leading to the activation of NF-κB, JNK, and p38 pathways. To explore the involvement of downstream proteins in TRAF6 condensates, we overexpressed mCherry-tagged TAK1, TAB3, IKKα, IKKβ, and NEMO proteins alone or together with EGFP-TRAF6 in HeLa cells. The results showed that when expressed alone, TAK1, TAB1, TAB3, IKKα, IKKβ, and NEMO were diffusely distributed in cytosol (Figure EV4A). However, when co-expressed with EGFP-TRAF6 in HeLa cells, these proteins all co-localized with EGFP-TRAF6 condensates (Figure 4A). In addition, after LPS stimulation, phosphorylated TAK1 (p-TAK1) exhibited co-localization with TRAF6 in condensates in Raw264.7^EGFP-TRAF6^ cells (Figure 4B), indicating that TRAF6 condensation promotes TAK1 phosphorylation and activation. Analogous to TAK1, NEMO also co-localized with TRAF6 condensates in Raw264.7^EGFP-TRAF6^ cells once stimulated (Figure 4C). Subsequently, our in vitro reconstitution data using purified TRAF6 and TAB1 peptides showed that TAB1 peptides were present in TRAF6 condensates beyond a concentration of 5 μM (Figure 4D). In vitro phase separation of TRAF6 and NEMO exhibited that the fluorescent signal of NEMO increased in TRAF6 condensates with increasing concentrations (Figure 4E). However, NEMO protein alone could not generate condensates in vitro (Figure 4B). These results collectively demonstrate that TRAF6 phase separation can recruit downstream proteins into condensates. Recently, it was reported that NEMO undergoes liquid phase separation induced by polyubiquitin chains (Du *et al*, 2022). In the process of IL-1β-induced NF-κB activation, TRAF6 is upstream of NEMO and responsible for polyubiquitin chain production, thus we speculate that both TRAF6 itself and its product - polyubiquitin chains promote NEMO phase separation. TRAF6 may function as the driving force to initiate phase separation of downstream signaling molecules.

**Figure 4.**
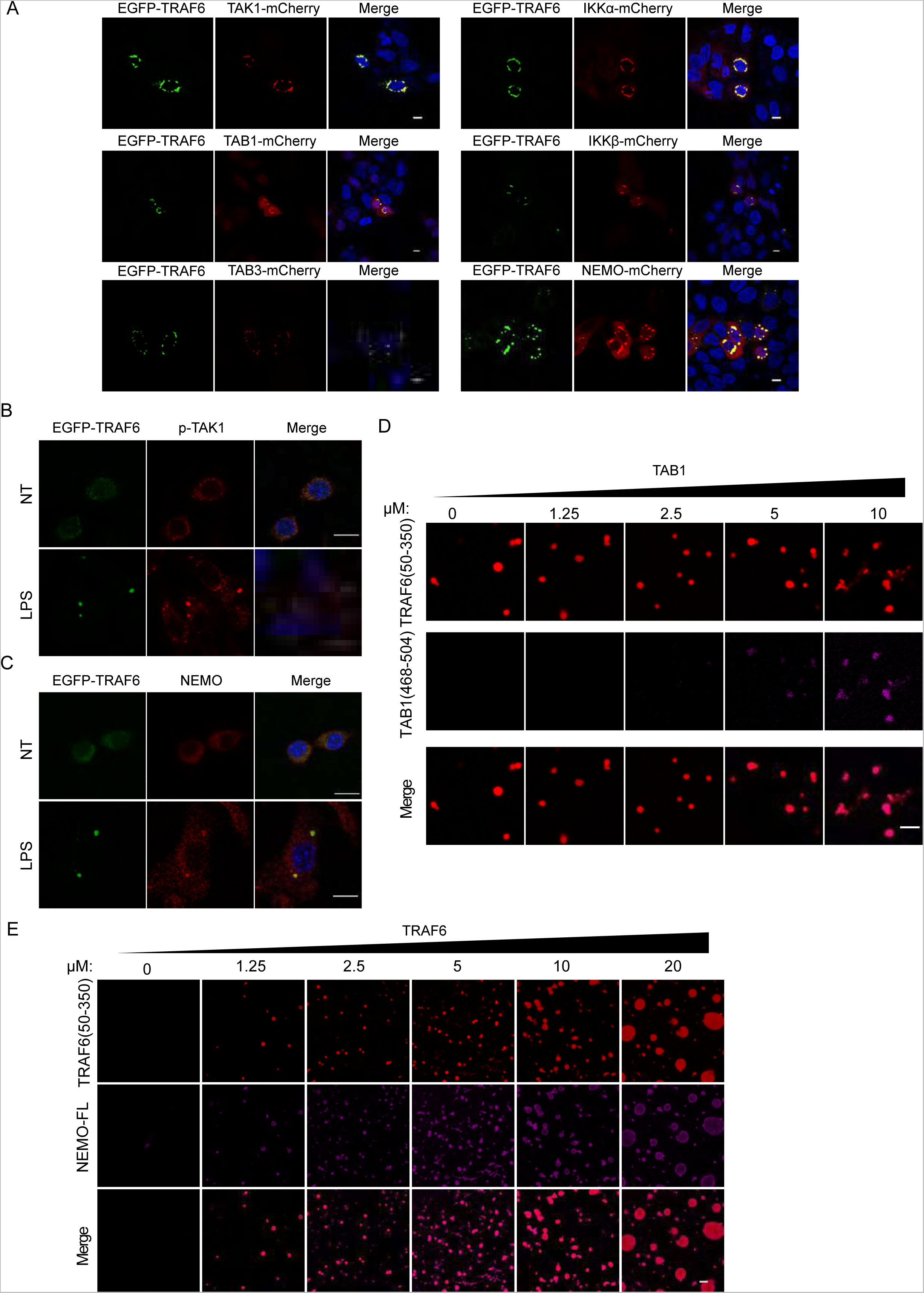
TRAF6 condensates recruit downstream proteins. A. Representative images of EGFP-TRAF6 co-expressed with TAK1-mCherry, TAB1-mCherry, TAB3-mCherry, IKKα-mCherry, IKKβ-mCherry and NEMO-mCherry separately. B,C. Representative colocalization images of Raw264.7^EGFP-TRAF6^ cells stimulated with or without LPS for 12 hours. Phosphorylated TAK1 (B) and NEMO (C) were separately stained with indicated antibodies. D. Representative colocalization images of TRAF6 and TAB1 in vitro at indicated concentrations. E. Representative colocalization images of TRAF6 and NEMO in vitro at indicated concentrations. The concentration of TRAF6 in (d) is 10 μM. The concentration of NEMO in E is 20 μM. Data information: Scale bars, 10 μm in (A-E).

### TRAF6 phase separation functions to activate inflammatory signaling

Next, to better understand the function and regulation of TRAF6 LLPS, an artificial LLPS inducing construct was made by replacing the coiled-coil domain with the FUS 1-162 amino acid domain (FUS(1-162)) (Figure 5A), which is capable of inducing phase separation (Patel *et al*, 2015; Burke *et al*, 2015). When overexpressed alone, FUS(1-162) formed few droplets in the nucleus (Figure 5F). However, TRAF6(50-286)-FUS(1-162) fusion protein produced many condensates in the cytosol (Figure 5H), suggesting that the coiled-coil domain and FUS(1-162) has a similar ability to induce TRAF6 LLPS. Moreover, in the NF-κB luciferase reporter assays, TRAF6(50-286) had no activation of NF-κB (Figure 5B and C). While TRAF6(50-349), which retained the ability to phase separate, had comparable activation of NF-κB to that of wild-type TRAF6 (Figure 5 B and C), suggesting that the coiled-coil domain is crucial for effective NF-κB activation. FUS(1-162) alone could not activate NF-κB signaling. However, TRAF6(50-286)-FUS(1-162) had strong activation of NF-κB, comparable to that of TRAF6(50-349) (Figure 5 B and C). These data indicate that LLPS of TRAF6 strongly promotes the activation NF-κB pathway.

**Figure 5.**
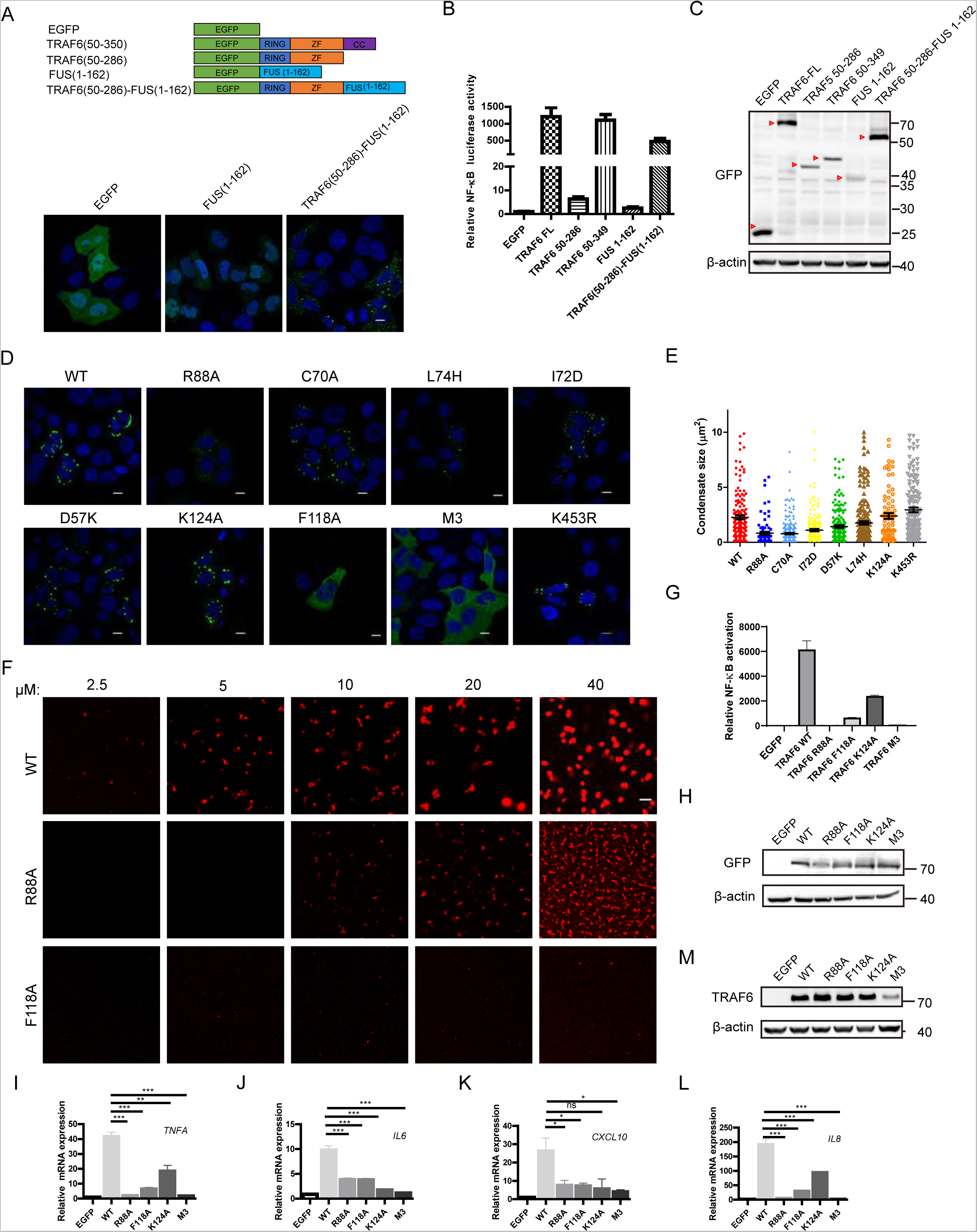
TRAF6 phase separates to activate downstream NF-κB signaling. A. Representative images of EGFP, EGFP-FUS(1-162) and EGFsP-TRAF6(50-286)-FUS(1-162) overexpressed in HeLa cells. B. NF-κB reporter results of TRAF6 truncations, deletions and FUS fusions in HEK293T cells. C. Western blotting of protein expressions in (B). D. Representative images of wild type EGFP-TRAF6 (WT) and its mutations, R88A, C70A, L74H, I72D, D57K, K124A, F118A, K453R and M3 overexpressed in HeLa cells. E. Statistic analysis of condensate sizes in (D). F. Representative images of in vitro phase separation of WT TRAF6, R88A TRAF6 and F118A TRAF6 proteins at indicated concentrations. G. NF-κB reporter results of TRAF6 wt and mutants in HEK293T cells. H. Western blotting of protein expressions in (G). I-L. The relative mRNA expression level of *CXCL10, TNFA, IL6*, *IL8* in the HeLa cells after overexpression at 24h. M. Western blotting of TRAF6 and mutants overexpression in HeLa cells in (I-L). Data information: Scale bars, 10 μm in (A, D and F). All data are representative of at least three independent experiments with similar results and the data in (B, E, G and I-L) are expressed as mean±SEM of technical replicates. *P < 0.05, **P < 0.01 and ***P < 0.001 by Student’s t test; ns, not significant.

To further explore the regulation mechanism of TRAF6 phase separation, a series of TRAF6 mutations were overexpressed in HeLa cells, including D57K, C70A, I72D, L74H, R88A, F118A, K124A, K453R and M3(I307A, L310A, I342A, L345A) (Yin *et al*, 2009; Hu *et al*, 2017). The D57K, I72D and L74H mutations disrupted TRAF6 interaction with Ubc13/UBE2N while the R88A mutant disrupted TRAF6 homodimerization. The C70A mutant impaired TRAF6 E3 ligase activity and the F118A mutant impaired both its homodimerization and polyubiquitin synthesis. The K124A mutant effects the self-ubiquitination of TRAF6. M3 mutant disrupted the interactions between coiled coils. Imaging and statistical results showed that the R88A, C70A, L74H, I72D, and D57K mutations significantly reduced the sizes of TRAF6 condensates (Figure 5C and D), while the K124A and K453R mutation had no significant effects on condensate sizes (Figure 5 C and D). Besides, the F118A and M3 mutants almost abolished TRAF6 condensate formation (Figure 5D and E). These data indicate that the dimerization, oligomerization, and interaction with E2 are crucial for regulation of its phase separation. When EGFP-TRAF6 proteins were overexpressed in Ubc13/UBE2N or Uev1A/UBE2V1 knockdown HeLa cell lines (Figure EV5A), the numbers and sizes of TRAF6 condensates were significantly reduced (Figure EV5B). C25-140, a small molecular inhibitor of TRAF6 - Ubc13/UBE2N interaction, also strongly inhibited TRAF6 droplets generation (Figure EV5C). These data all suggest that TRAF6 phase separation in cells is modulated by self-interaction, interaction with E2s and polyubiquitin synthesis. Whereafter, we performed in vitro LLPS experiments with mutant proteins. Consistent with results in cells, higher concentrations were required for the TRAF6-R88A protein to form condensates compared to wild-type protein (Figure 5C), and the condensates formed by TRAF6-R88A protein were much smaller at the same concentrations (Figure 5F). Remarkably, TRAF6-F118A protein rarely formed condensates even at a high concentration of 40 μM (Figure 5F). These in vitro data consistently underlie that both self-multivalent interactions and polyubiquitin chain synthesis ability of TRAF6 regulate its LLPS.

The NF-κB luciferase assay results demonstrated that TRAF6-R88A, TRAF6-F118A, TRAF6-M3 proteins had no activation of NF-κB signaling (Figure 5G and H). Consistently, TRAF6 mutants without the ability to phase separate lost the ability to induce expression of cytokines, including *TNFA*, *CXCL10* and *IL6* (Figure 5I-L). In summary, we conclude that TRAF6 phase separation is fundamental for the activation of inflammatory signaling.

In brief, we proposed a model for TRAF6 initiated phase separation (Figure 6). When stimulated by TLRs or IL-1R agonists, TRAF6 will sense the upstream signals and phase separate into gel-like condensates, recruiting ubiquitin synthesis components, TAB1, TAK1 and NEMO, leading to the phosphorylation and ubiquitination of a series proteins, finally activation of inflammatory signaling.

**Figure 6.**
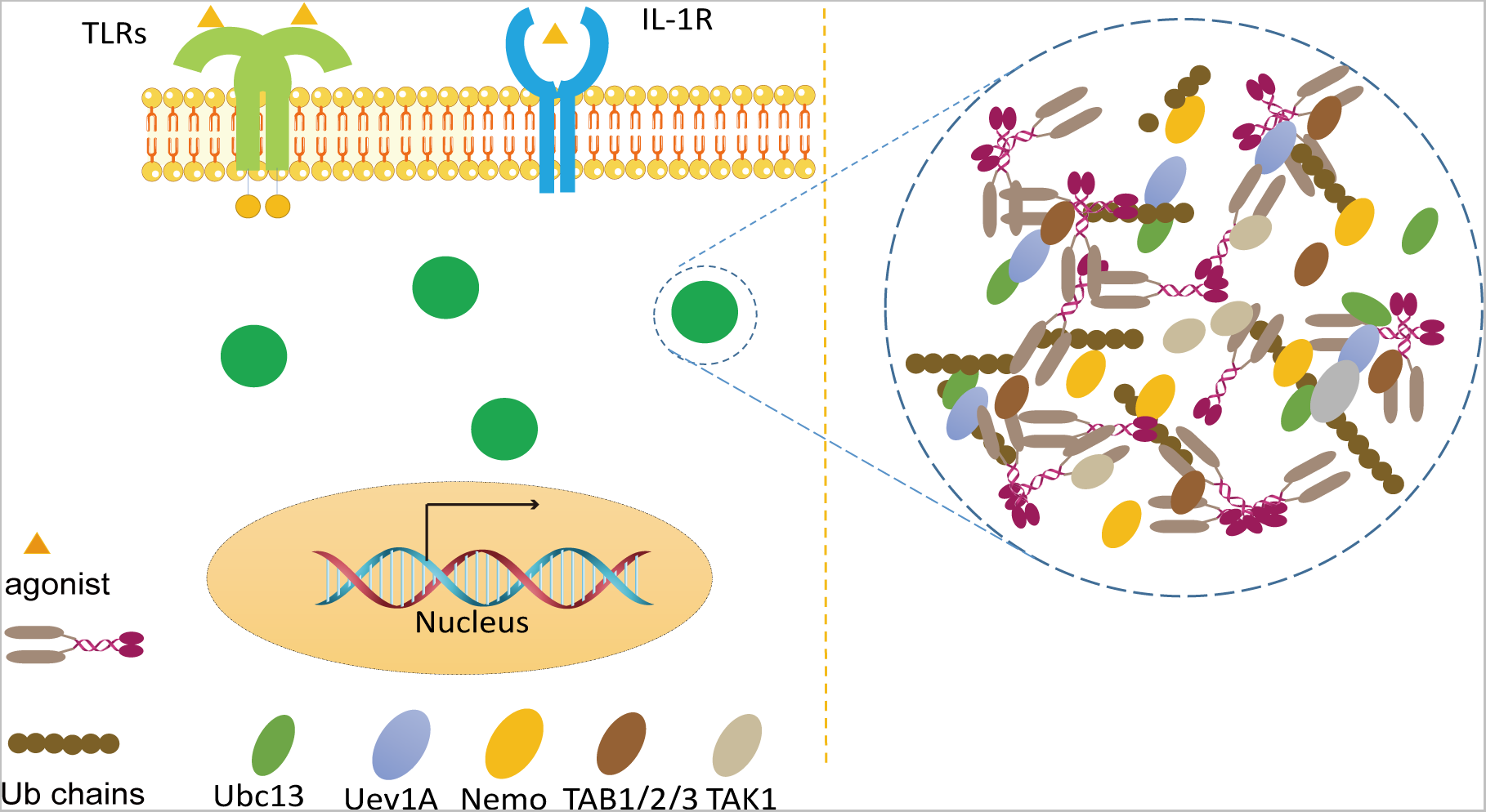
Schematic diagram of the model of TRAF6-initiated phase separation.

## Discussion

Our results indicate that TRAF6 complex undergo phase separation to form membranless condensates to initiate downstream activation of inflammatory signaling. Although LLPS has been found involving multiple processes, the role of LLPS in transduction of signals form membrane is only emerging (Li *et al*, 2012; Banjade & Rosen, 2014; Su *et al*, 2016; Bonello *et al*, 2023). We prove that the multivalent interaction of TRAF6 leads to phase separation during PAMP stimulation from the environment. TRAF6 condensation may serve as a rapid sensor to combat pathogens invading. This quick and efficient way to transmit danger signals is important for protection from bacteria and parasites. Additionally, in the process of Gram-negative bacteria infection, the ALPK1-TIFA-TRAF6 axis is activated by the metabolite ADP-β-d-manno-heptose (Gaudet et al, 2015; Zhou et al, 2018). TIFA can form punta structure with TRAF6 (Gaudet et al, 2015), indicating TRAF6 may undergo LLPS in defense against Gram-negative bacteria.

Its RING domain can form both homo- and hetero-dimers (Yin *et al*, 2009; Middleton *et al*, 2017). This domain also interacts with Ubc13/UBE2N and ubiquitin to facilitate ubiquitin transfer. The coiled-coil domain of TRAF6 can induce its oligomerization, ensuring efficient assembly of long polyubiquitin chains (Hu *et al*, 2017). Previous model from structure view only assume TRAF6 forms dimer, trimer or oligomers(Ye *et al*, 2002;Yin *et al*, 2009; Ferrao *et al*, 2012; Middleton *et al*, 2017; Fu *et al*, 2018). In our study, we further prove that the combination of these multiple interactions leads to a more complex and higher-level of assembly, i.e., LLPS. At endogenous level, the protein level of TRAF6 was much lower than that by overexpression, the phase separation of TRAF6 ensures rapid and efficient responses to pathogens or damages. As for how TRAF6 is activated, we speculate the upstream complex formed by MyD88/IRAK1/IRAK4 probably serve as a starting trigger to concentrate TRAF6.

The TRAF family proteins contain 7 TRAF proteins. TRAF 2-7 all have the RING domain, the zinc finger domain, and the coiled coil domain; TRAF1 lacks the RING domain (Zhu *et al*, 2018). Whereupon we overexpressed these EGFP tagged TRAF family proteins in HeLa cells, and the results showed that all TRAF family proteins could form condensate structures like TRAF6 (Figure EV6A). The data indicate that TRAF family proteins have the capability to phase separate and may function through phase separation. One recent study reports that by application of coiled-coils, the antibody is prevented from antigen sink and gains increased anti-tumor efficacy (Trang *et al*, 2019). In the future, new drugs can be developed by leveraging the ability of TRAF family proteins to induce phase separation.

Ubiquitination is a crucial phenomenon that regulates diverse aspects of immune functions (Hu & Sun, 2016). Ubiquitin play a complex role in LLPS (Dao *et al*, 2018; Yasuda *et al*, 2020; Dao *et al*, 2022). In our investigation, we reveal that TRAF6 itself undergoes LLPS and recruits downstream proteins. Recently, Du et al reported that unanchored polyubiquitin chains have the ability to promote LLPS of NEMO (Du *et al*, 2022). Thus, TRAF6 together with the long polyubiquitin chains probably forms a “reaction factory” to facilitate phosphorylation and ubiquitination reactions via promoting proximity. The interactions between TRAF6 itself and downstream TAB1, TAK1, NEMO and IKKs at the atomic level call for deeper investigation by structure biologists and biophysicists. In several types of cancers, TRAF6 mRNA and protein level are elevated with increased NF-κB activation, such as lung cancers (Starczynowski *et al*, 2011), breast cancer (Rezaeian *et al*, 2017) and melanoma (Liu *et al*, 2012). Since high protein concentration can induce TRAF6 phase separates, the aberrant high levels of TRAF6 probably lead to auto-activation of TRAF6 and NF-κB signaling in cancer development.

In summary, our study adds to the puzzle of phase separation in innate immunity and provides a theoretical basis for the development of therapies targeting TRAF6 phase separation.

## Materials and Methods

### Reagents and Tools table

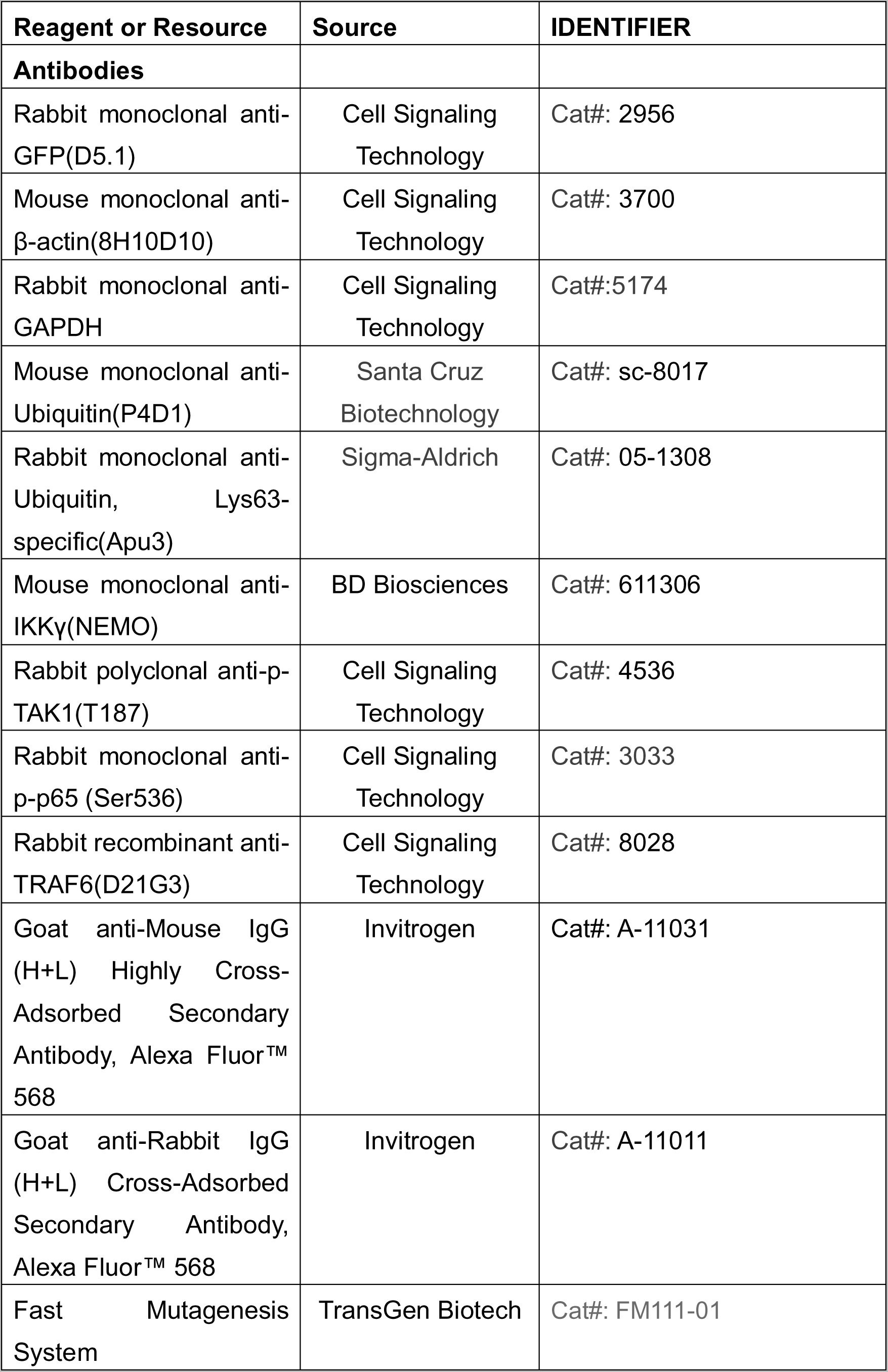
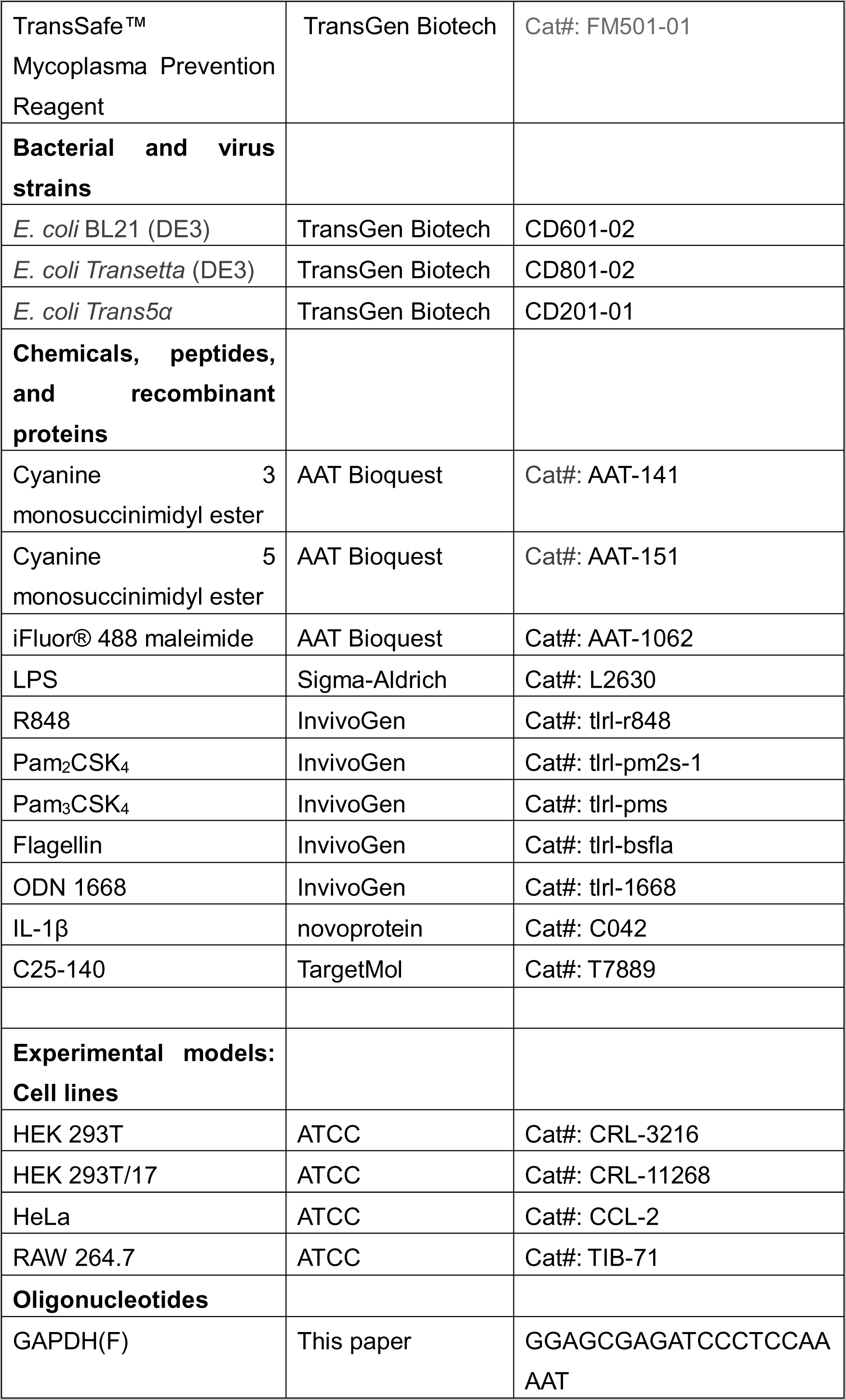
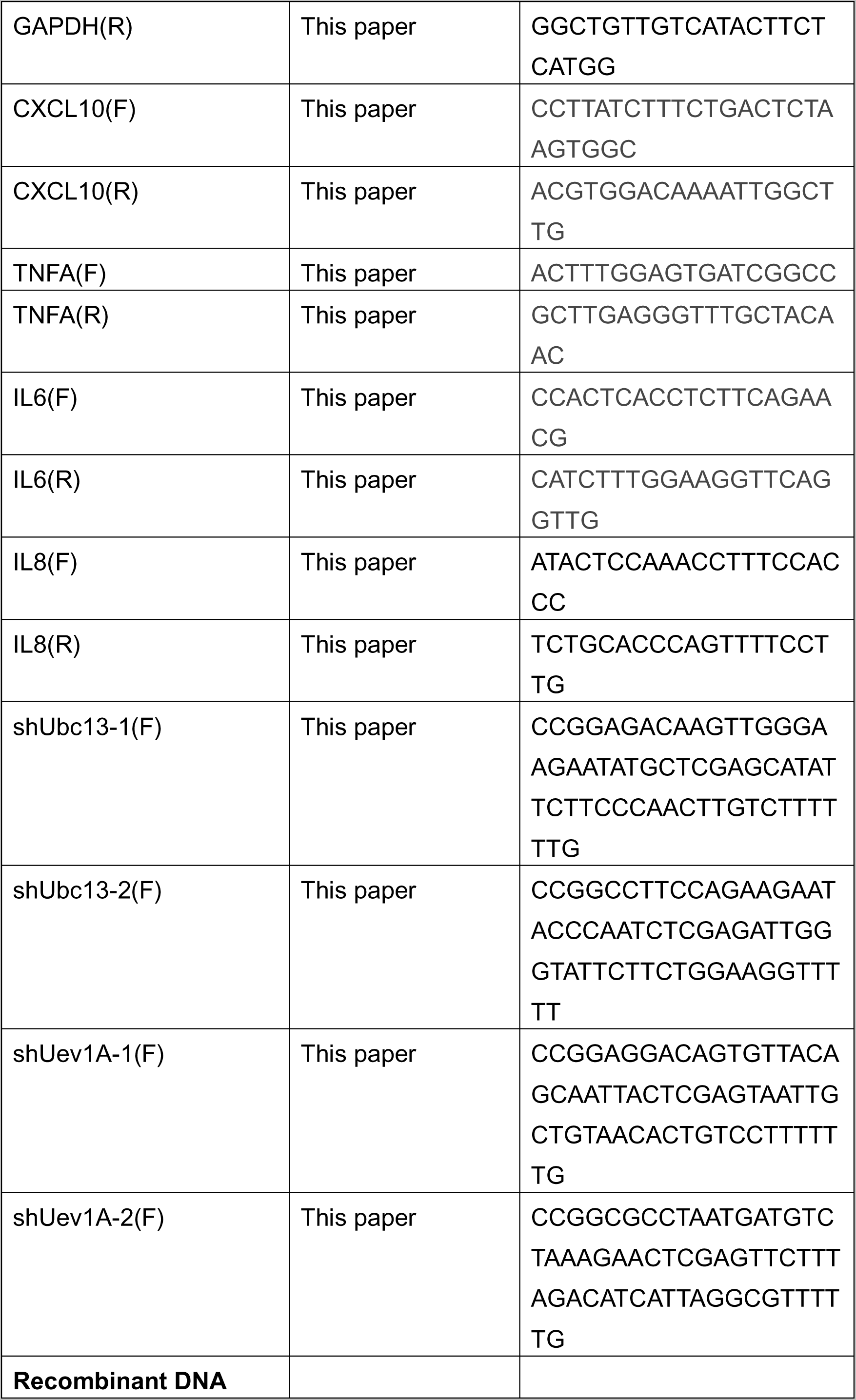
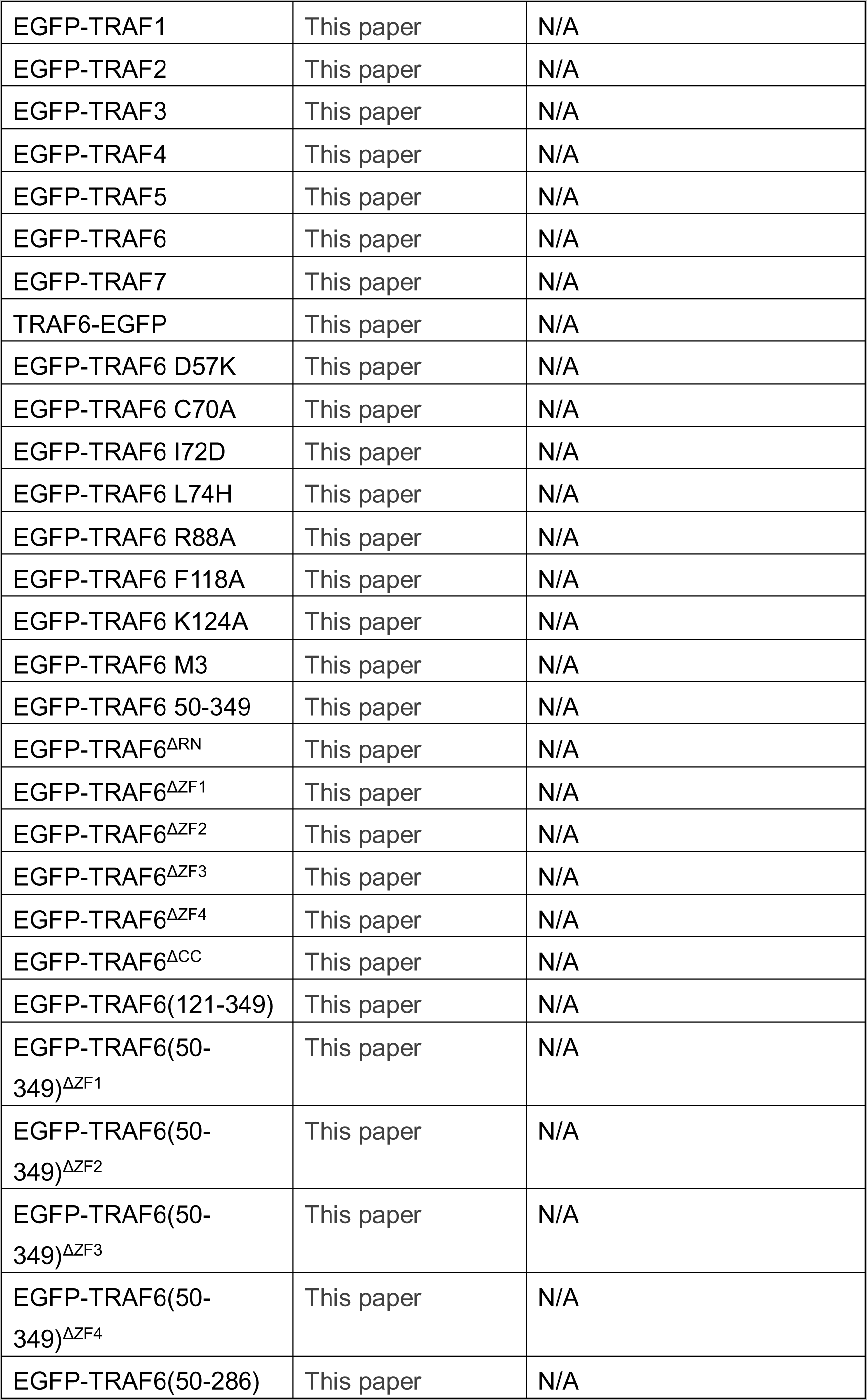
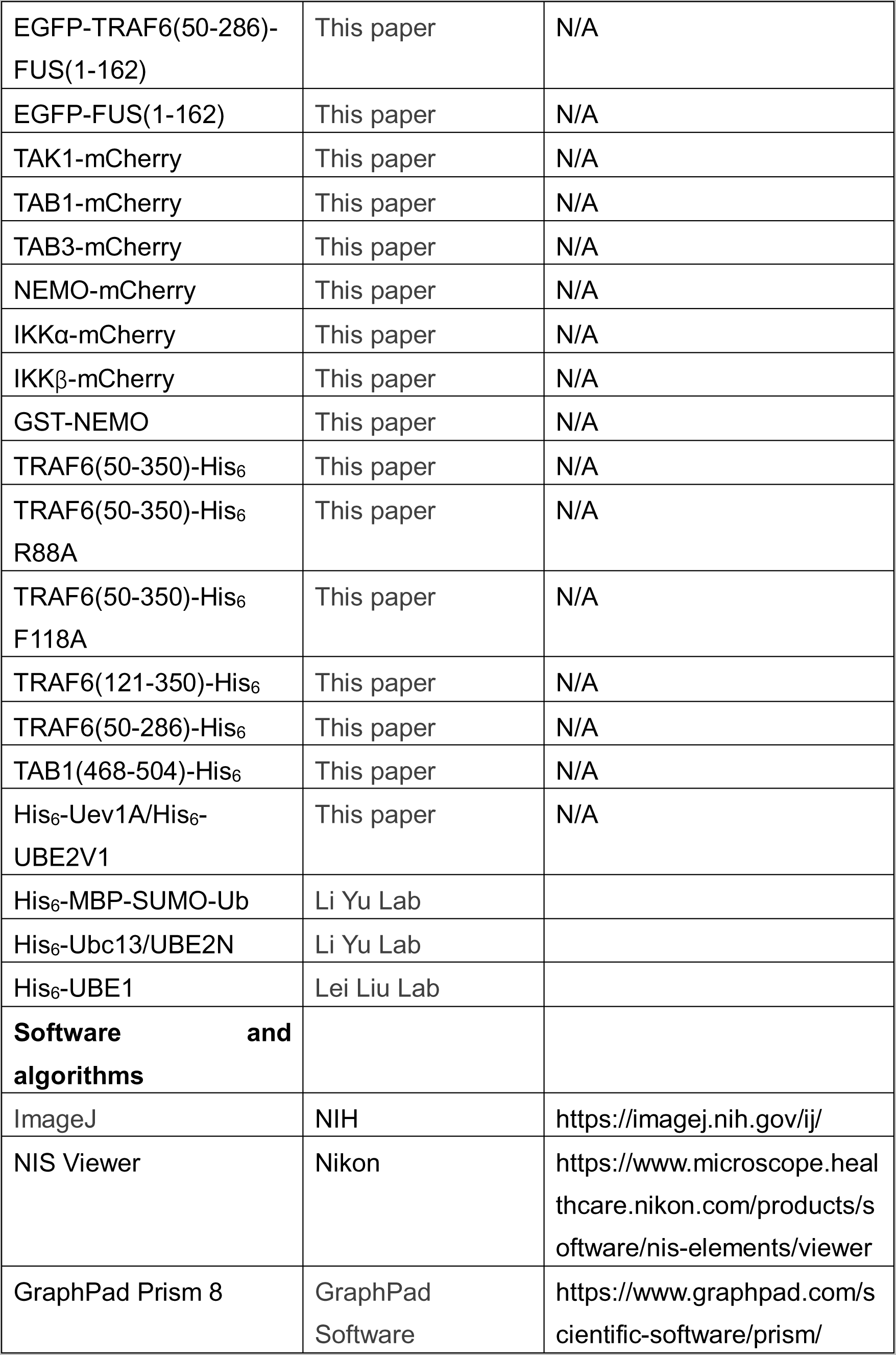

### Methods and Protocols

#### Cell Culture, treatment, and transfection

HEK 293T cells, HeLa cells and RAW 264.7 cells were cultured in DMEM (Invitrogen) ed with 10% FBS (Invitrogen), 100 U/ml penicillin, and 100 μg/ml streptomycin. All cells were maintained at 37°C in an atmosphere of 5% CO_2_. Cells were treated with TransSafe™ Mycoplasma Prevention Reagent (TransGen Biotech). HEK 293T cells and HeLa cells were transfected with plasmids using lipofectamine 3000 according to the manufacturer’s instruction (Invitrogen).

#### Plasmid construction

The human TRAF6 WT, variants and truncations were cloned into BglⅡ- HindⅢ sites of pEGFPC1 vectors through Gibson assembly. TAK1, TAB1, TAB2, IKKα, NEMO, IKKβ, p65, JNK, p38 were cloned into XhoⅠ-HindⅢ sites of pmCherryN1 vectors, part of the templates were kindly provided by Li Yulong lab from Peking University. The FUS template (525AA) was a gift from Li Pilong lab. FUS 1-162 is fused to the C-terminal of EGFP-TRAF6 50-286.Human NEMO and was cloned into BamHⅠ-EcoRⅠof pGEX-6p-1 vector and a HRV 3C protease recognition sequence was added between GST tag and target genes. All mutations were made using the Fast Mutagenesis System. Human TRAF6 (50-350), TRAF6 (50-350) R88A, TRAF6 (50-350) F118A, TRAF6 (121-350), TRAF6(50-286) and TAB1 (468-504) were cloned into Nde Ⅰ-BamHⅠsites of pET21a(+). 2 amino acids M-K was added to the N terminal of TAB1 (468-504).

His_6_-Ubc13 and His_6_-MBP-SUMO-Ub construct was provided by Prof. Yu Li and 3 amino acids S-C-G was added to the N terminal of ubiquitin protein after SUMO protease digestion. His_6_-UBE1 construct was donated by Prof. Lei Liu.

His_6_-Uev1A/UBE2V1 were cloned into NdeI-BamHI sites of pET28a. GFP-TRAF6 was cloned into pQCXIP plasmid at BglII-EcoRI sites.

#### Protein expression, purification and labeling

Recombinant TRAF6(50-350) proteins and other variants were expressed in the *E.coli* strain *Transseta* transformed with corresponding plasmids. All TRAF6 constructs were induced with 0.2 Mm IPTG, 0.1 mM ZnCl_2_ at 16 °C for 16hr. After induction with 0.2mM IPTG at 18 °C for 20 hours, the bacteria were collected and lysed in the lysis buffer containing 25mM Tris-HCl, pH 8.0, 150mM NaCl, and 1mM PMSF with high pressure. The lysates supernatant was incubated with Chelating Sepharose pretreated with 0.2 M NiSO_4_ and washed with an elution buffer containing 25mM Tris-HCl, pH 8.0, 150mM NaCl and Imidazole ranging from 50mM to 300mM. The proteins were then purified on AKTA equipped with a Source Q column. The purity of protein fractions was detected with SDS-PAGE. The protein fractions with high purity were concentrated with a centrifugal concentrator (Millipore). TAB1 (468-504) were expressed in the *E.coli* strain *Transseta*(DE3) in a process similar to that of TRAF6(50-350).

GST-NEMO was expressed in the *E.coli* strain *Transseta*(DE3) transformed with the pGEX-6p-1 plasmid encoding the indicated protein. After induction with 0.5mM IPTG at 18 °C for 20 hours, the bacteria were collected and lysed in a lysis buffer containing 25mM Tris-HCl, pH 8.0, 150mM NaCl, and 1mM PMSF with high pressure. After centrifugation, lysate supernatant was incubated with Glutathione Sepharose 4B resin (GE Healthcare) followed by loading onto gravity flow columns, washing with lysis buffer and the GST tag was cleaved on column with homemade GST-HRV 3C protease at 4 °C overnight. Cleaved protein was concentrated with a 30 kDa MWCO, centrifugal concentrator (Millipore).

His_6_-MBP-SUMO-Ub was expressed in the *E.coli* strain BL21(DE3) with 0.5mM IPTG induction. Similar to TRAF6(50-350) purification, the eluted proteins were cleavage with sumo protease overnight in a dialysis buffer containing 25mM Tris-HCl, pH 8.0, 20mM NaCl and purified on AKTA equipped with a Source Q column. The protein fractions were collected and concentrated using a 3 kDa centrifugal concentrator.

Ubc13, Ube2v1 and UBE1 were expressed in the *E.coli* strain BL21(DE3) with 0.5mM IPTG induction. The purification procedure was the same with TRAF6(50-350). For Ubc13 and Ube2v1, the eluted proteins were loaded onto a superdex75 column equilibriumed with 25mM Tris-HCl, pH 8.0, 150mM NaCl, 2mM DTT. For UBE1, the eluted proteins were loaded onto a superdex200 column equilibrated with 25mM Tris-HCl, pH 8.0, 150mM NaCl, 2mM DTT.

Recombinant TRAF6(50-350) and other variants were labeled with Cyanine 3 (AAT Bioquest), UBE2N, NEMO and TAB1(468-504) were labeled with Cyanine 5 (AAT Bioquest), Ub was labeled with iFluor® 488(AAT Bioquest) according to the manufacturer’s instruction.

#### Cell lines construction

HEK 293T/17 cells were transfected with retroviral transfer plasmids pQCXIP-EGFP-TRAF6 plus packaging plasmids pVSVG and pHIT and the virus particles were harvested 48hr after transfection, filtered through a 0.45μm membrane filter and stored in −80°C. RAW 264.7^EGFP-TRAF6^ stable cells were generated by infecting the RAW 264.7 with harvested retrovirus in the presence of 1μg/ml polybrene. After 48hr of culture, transduced cells were selected with 2μM puromycin. The positive cells were verified by Western blotting and RT-qPCR analyses. LPS (100 ng/mL), IL-1β(100 ng/mL), Pam_2_CSK_4_ (100 ng/mL), Pam_3_CSK_4_ (100 ng/mL), flagellin (100 ng/mL), R848 (2 μg/mL) and ODN (1μM) were added to the DMEM without FBS for the indicated time.

pLKO.1 Ubc13/UBE2N-shRNA(TRC_ID: TRCN0000368937, TRCN0000007216) and pLKO.1 Uev1A/UBE2V1-shRNA plasmids(TRC_ID: TRCN0000033707, TRCN0000033708) were obtained to make stable knockdown cell lines from Center of Biomedical Analysis, Tsinghua University. HEK 293T/17 cells were transfected with the indicated lentivirus plasmids together with the packing plasmids psPAX2 and pMD2.G. The virus package, infection of HeLa and puromycin selection were similar to the generation of RAW 264.7^EGFP-TRAF6^ stable cells. The shRNA sequences used here are listed in Table 1. The knockdown cell lines were transfected with EGFP-TRAF6, then treated with 30μM or 60 μM C25-140 for 12hr or 24hr.

**Table 1.**
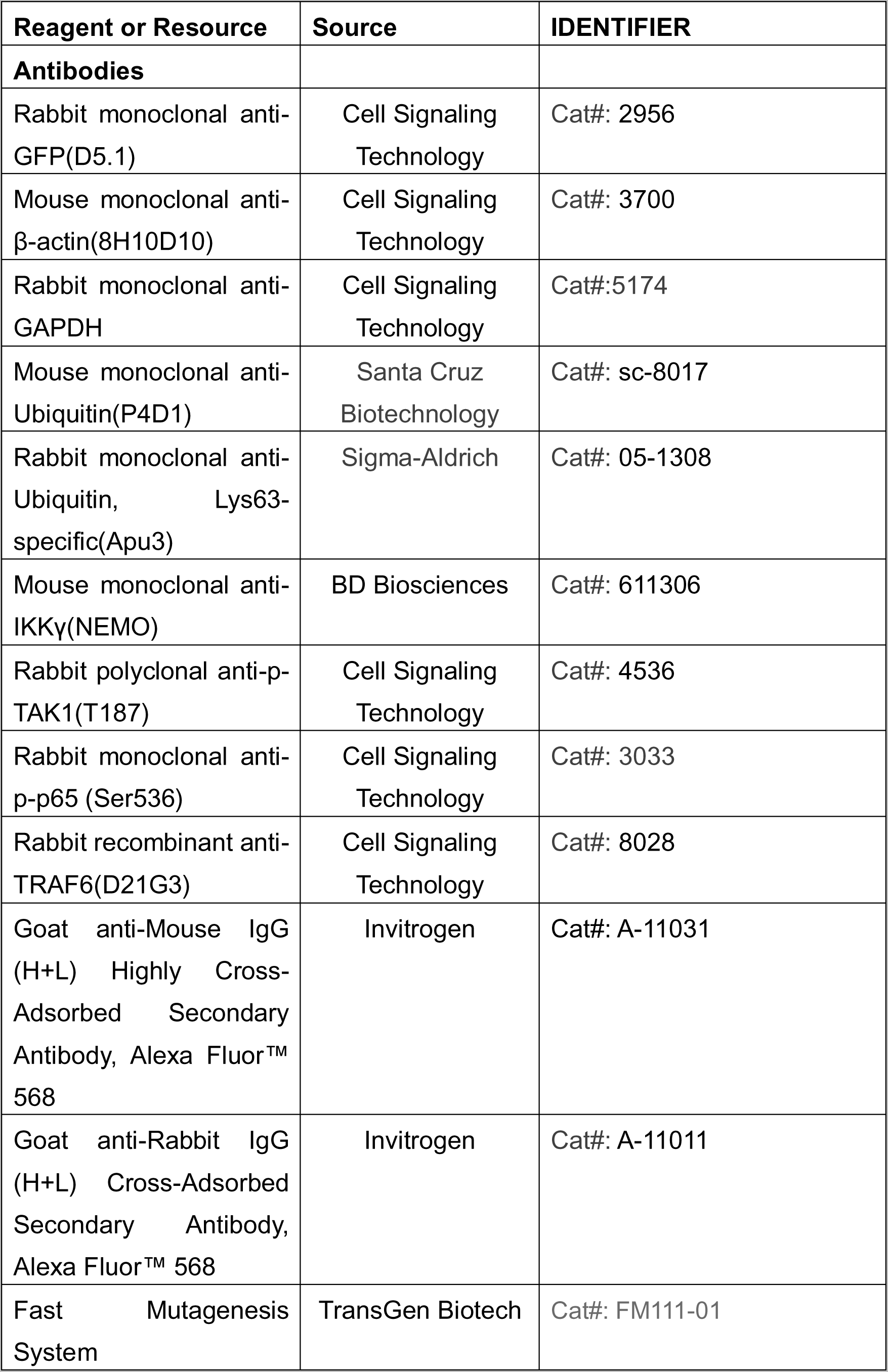
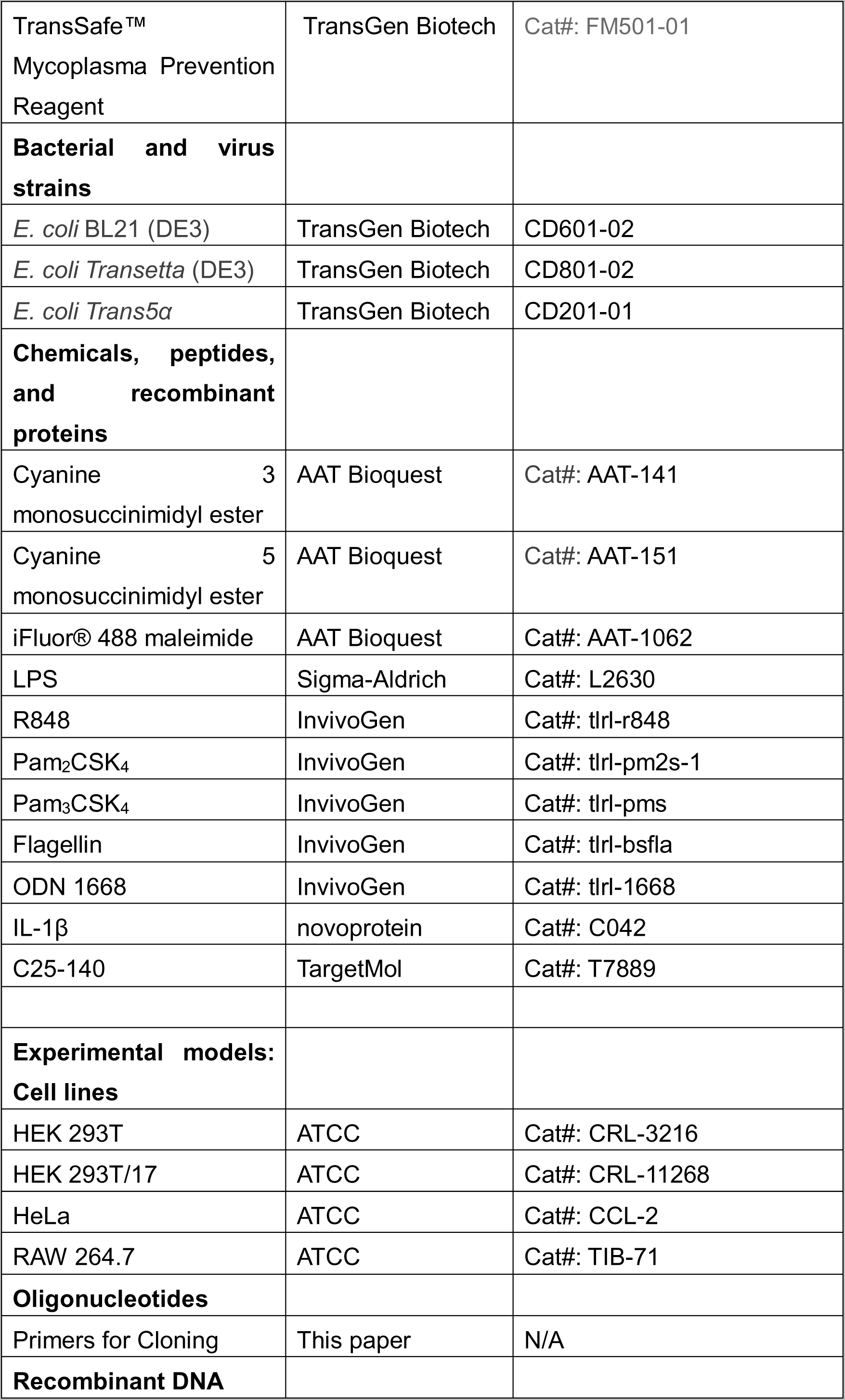
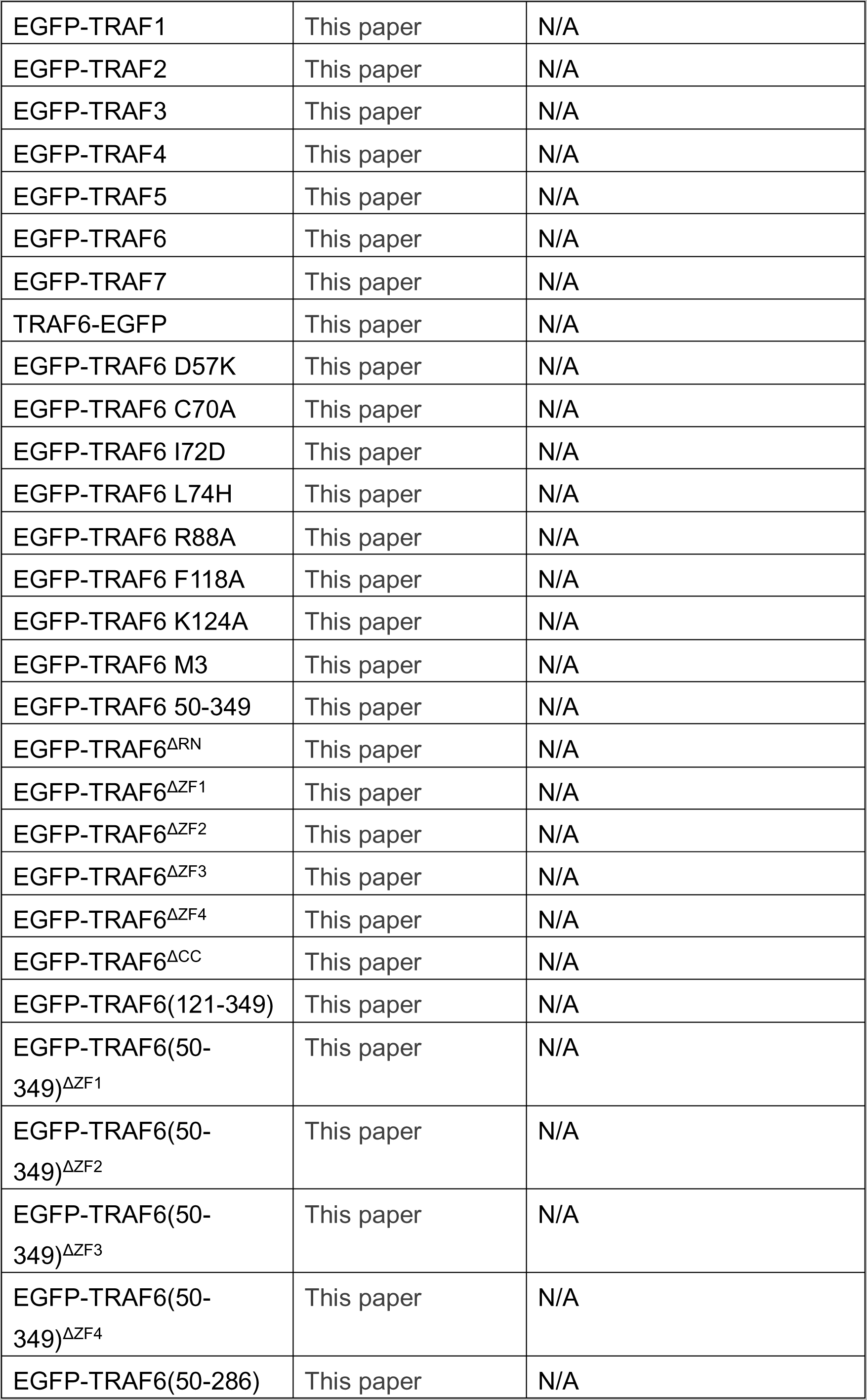
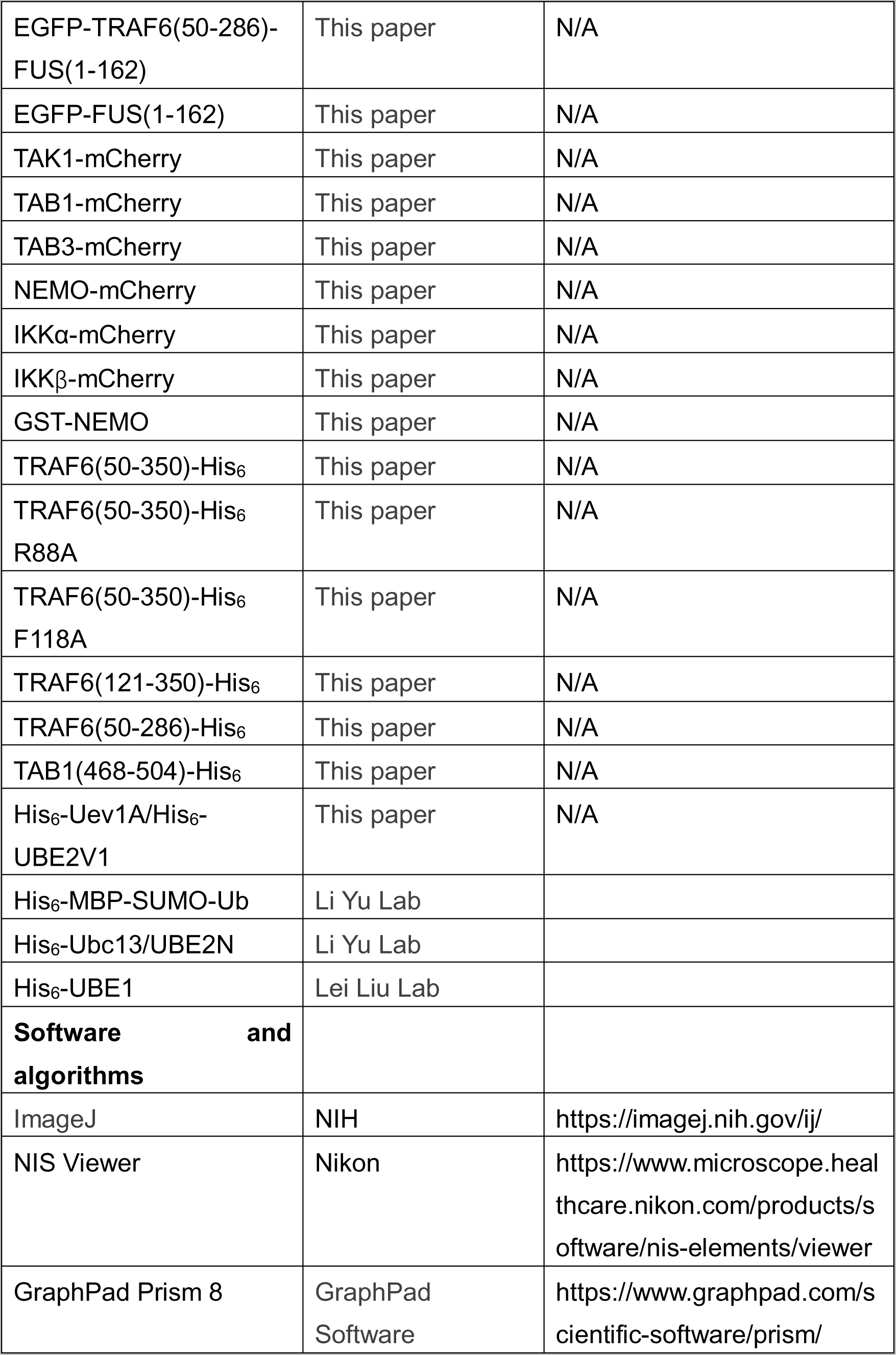
Materials.

**Table 2.**
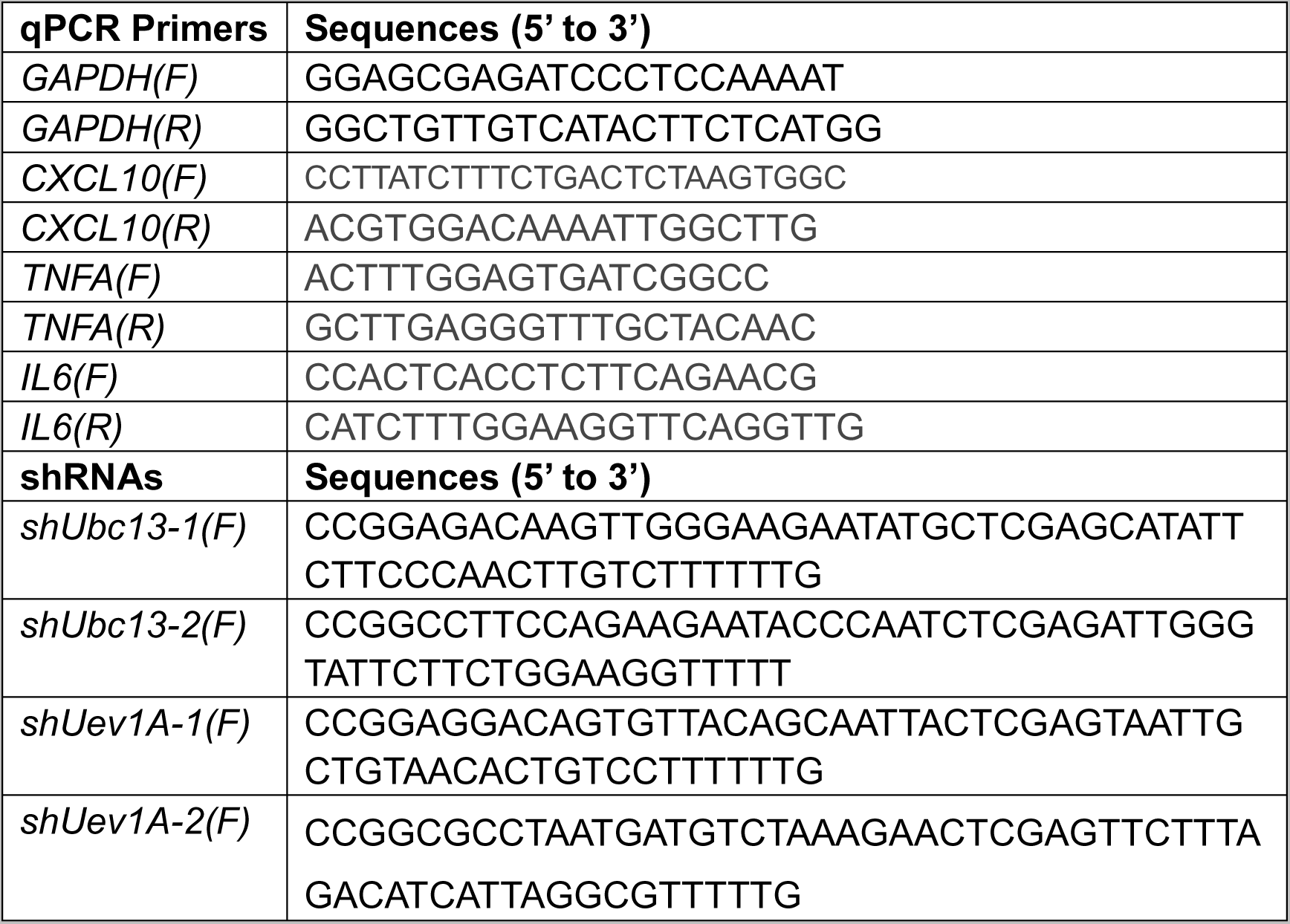
Primers.

#### In vitro phase separation

The purified TRAF6(50-350) and derivatives, NEMO, TAB1(468-504) (2% fluorescence labelled) were diluted to indicated concentration in a buffer containing 40mM Tris–HCl, pH 7.5, 5mM MgCl2, 150mM NaCl, 10% glycerol, 1mM DTT. In the ubiquitination system, 2mM ATP, 50μM Ub, 0.1μM UBE1, 1μM Ubc13, 1μM Uev1A were added to the mixtures. Mixtures were transferred to 384 well glass bottle plate(cellvis) and images were captured by confocal microscopy.

#### Sedimentation assay

After in vitro ubiquitination assay, TRAF6 droplets were spun down at 12,000 g for 10 min. The amount of ubiquitin or poly-ubiquitin chain was analyzed by western blot in the supernatant and pellet.

#### Western blot analysis

Cell lysates or in vitro reaction system were boiled in 1× Laemmli sample buffer for 5min, and then electrophoresed on proper condition. Proteins were transferred to 0.45 μm polyvinylidene difluoride (PVDF) membranes followed by 5% non-fat milk blocking. The first antibodies were all acquired commercially as previously mentioned. The blots were detected using HRP-conjugated secondary antibodies (Huabio) and ECL system. The blots were then imaged on an iBright 1000 Imaging System (Invitrogen) and analyzed by ImageJ.

#### Immunofluorescence staining

HeLa or RAW 264.7^EGFP-TRAF6^ cells were seeded on 20mm glass cover (NEST) at 8×10^4^ cells/well in a 12-well tissue-culture plate overnight. HeLa cells were transfected with plasmids using lipofectamine 3000 and RAW 264.7^EGFP-TRAF6^ cells were treated with TLR agonists. 24hr after transfection or 12hr after stimulation, cells were fixed with 4% paraformaldehyde in PBS at room temperature for 15 min. Cells were then permeabilized by 0.1%Triton-X100 in PBS for 15min. Fixed cells were blocked by 5% BSA in PBS for 1 hour followed by incubation with indicated antibodies in PBS containing 5% BSA at room temperature for 3hr. After 3 times of 5 in washes with PBS, cells were incubated with secondary antibody and DAPI on the shaker in light-free condition for 1hr. After 3 times of 5 in washes with PBS, all the processed slides were mounted in antifade mounting medium.

#### Live cell imaging and the FRAP assay

HeLa were grown on chambered cover glass (cellvis) to an appropriate density. 24hr after transfection, the droplets in EGFP-TRAF6 overexpressed cells were captured. During the imaging, cells were placed in a 37°C live-cell-imaging chamber supplied with 5% CO_2_. TRAF6 droplets were photobleached to 0-30% florescent intensity level of initial state. Time-lapse images were acquired over a 20min time course after bleaching with 20-second interval. For in vitro FRAP, the fluorescence was bleached with the corresponding laser and time-lapse images were acquired over a 10min time course after bleaching with 10-second interval.

#### Acquisition and Analyses of Images

All the confocal microscopy data was acquired from Nikon A1(R) HD25 equipped with 100×oil objective. Images were processed by NIS Viewer or ImageJ. Fluorescence intensities of regions of interest (ROIs) were corrected by unbleached control regions and then normalized to pre-bleached intensities of the ROIs.

#### Luciferase assay

4×10^5^ HEK 293T cells per well were seeded in the 12-well tissue-culture plate and were cultured overnight. HEK 293T cells were then transfected in triplicate with test plasmids plus 100ng NFKB Luciferase reporter construct. 24 hr after transfection, cells were lysed with Passive Lysis Buffer (Promega) and the luciferase activity is recorded by immediately after the substrate (Promega) is added.

#### qRT-PCR analysis

HeLa cells were seeded in 12-well plates and treated as indicated in the figure legends. Total RNA was extracted using the RNAsimple Total RNA kit (TIANGEN), and then was reverse-transcribed using an iScript cDNA synthesis kit (Novoprotein). The cDNA targets were analyzed by qPCR on a Bio-Rad T100 thermal cycler. Primer sequences are listed in key resource table.

#### Quantification and statistical analysis

Statistical analysis of the data were done using Graphpad Prism 5 software. The one-way ANOVA with Tukey post hoc test was performed for multiple comparisons. P values less than 0.05 were considered to be of statistical significance. Statistical difference was shown as *P < 0.05, **P < 0.01, and ***P < 0.001. NS P > 0.05 represents no significant difference.

## Acknowledgments

This work was supported by the National Natural Science Foundation of China (no. 21825702), National Key R&D Program of China (no. 2017YFA0505200), Beijing Outstanding Young Scientist Program Grand No. BJJWZYJH01201910003013 (H.Y.) and the Chun Feng Program of Tsinghua University (no. 2020Z99CFY036). We thank Prof. Li Yu for donating the His-Ubc13/UBE2N and His-sumo-UB plasmids; Prof. Lei Liu for donating the His_6_-UBE1 plasmid. We also thank Prof. Pilong Li for donating the FUS plasmid. We are grateful to Professor Conggang Zhang and Dr Ying Zhang for helpful discussion. We would like to acknowledge the assistance of SLSTU-Nikon Biological Imaging Center for the assistance using the Nikon A1R Instrument.

The manuscript was edited with assistance from ChatGPT.

## Author contributions

J.W. and H. Y. conceived the project. J.W. and X. Z. designed and performed all the experiments. J.W., X. Z., and H. Y. wrote the manuscript.

## Conflict of interest

The authors declare no competing interests.

Supporting Information accompanies the manuscript on the XXX website http://www.

## Expanded View Figure legends

**Figure EV1.**
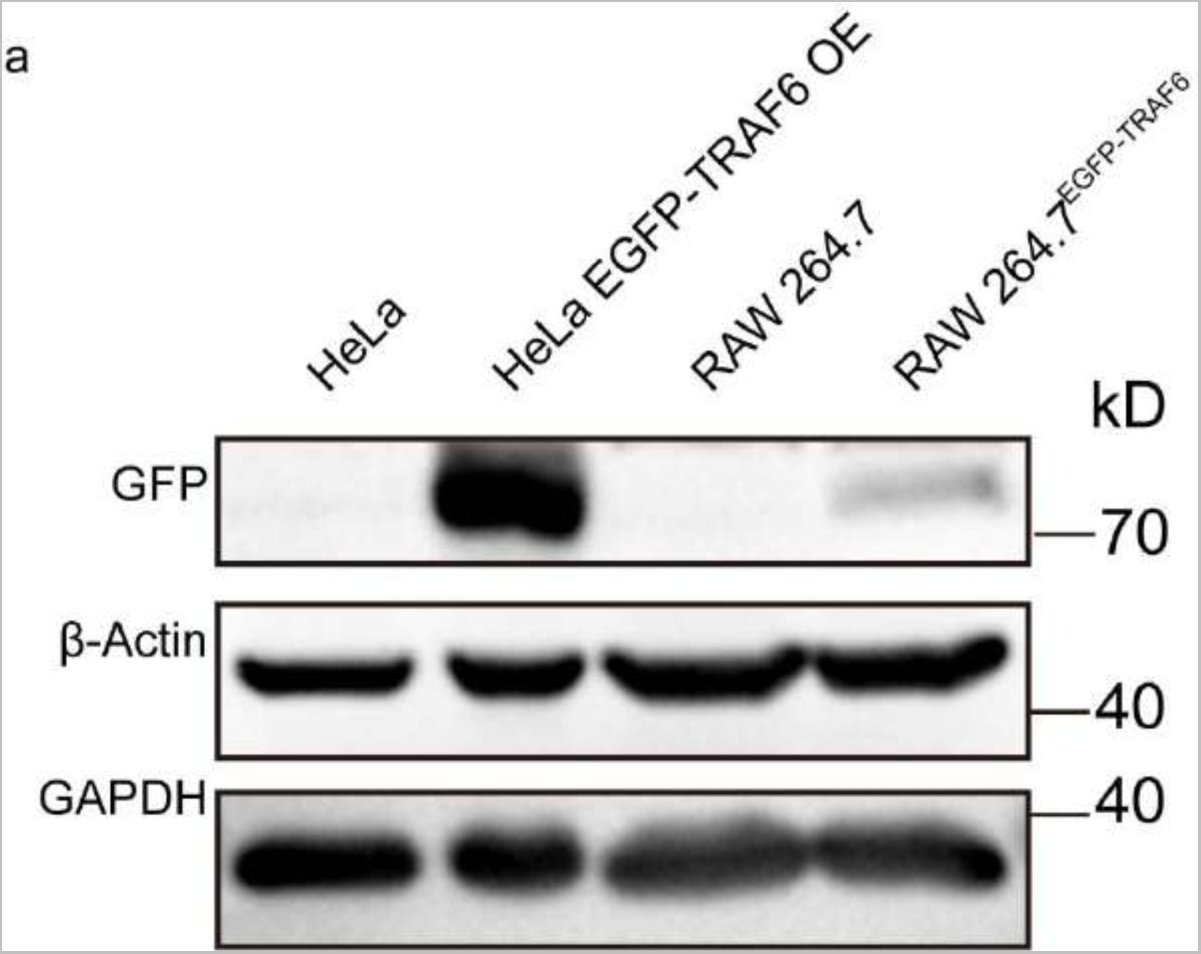
The TRAF6 expression level in cell lines. A. Comparision of EGFP-TRAF6 overexpressed in HeLa cells and EGFP-TRAF6 in Raw^EGFP-TRAF6^ cell lines.

**Figure EV2.**
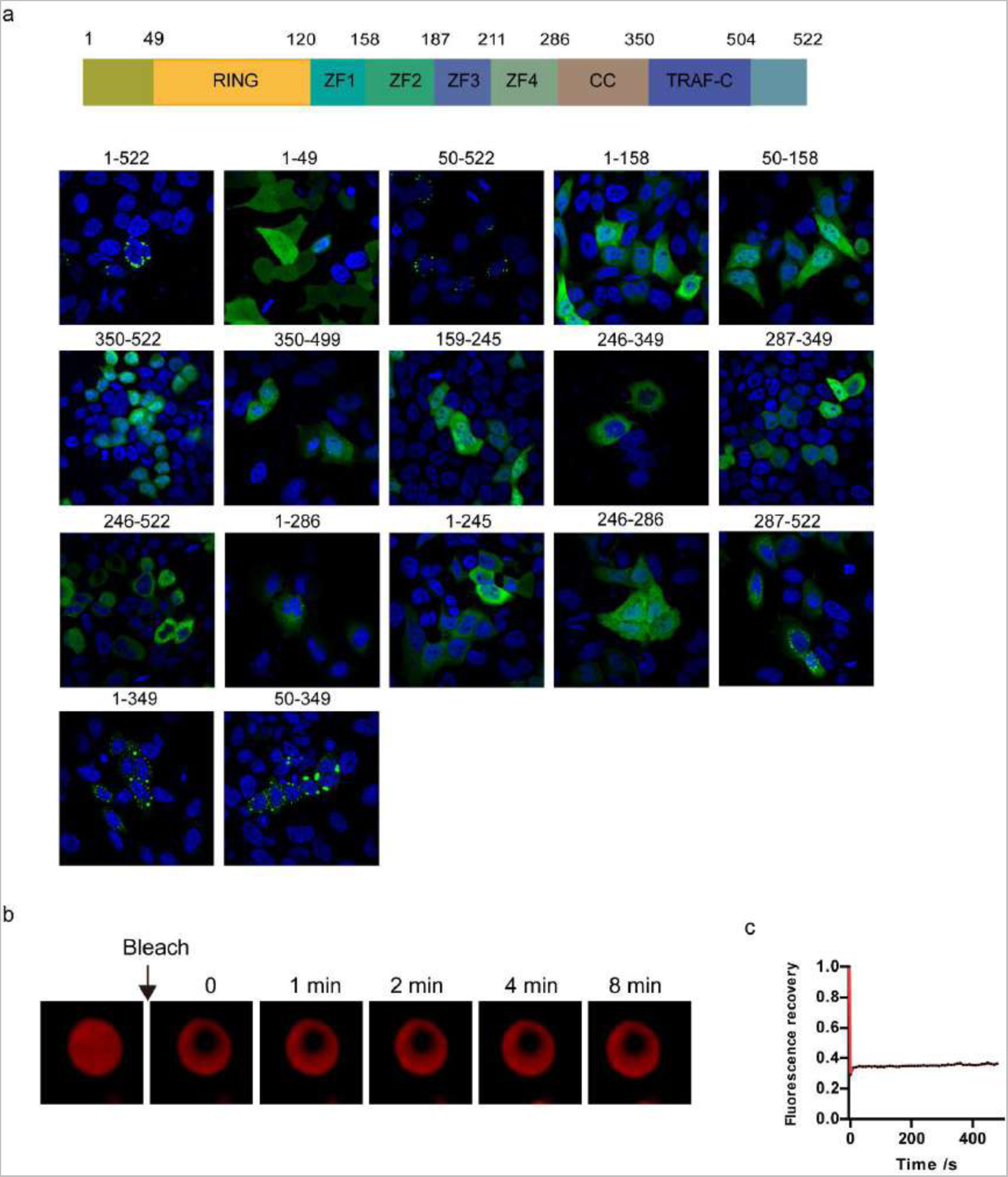
The phase separation ability of different TRAF6 truncations. A. Representative images of TRAF6 truncations in HeLa cells.
B. Representative FRAP images of TRAF6 condensates.
C. Statistic analysis of FRAP in (B)

**Figure EV3.**
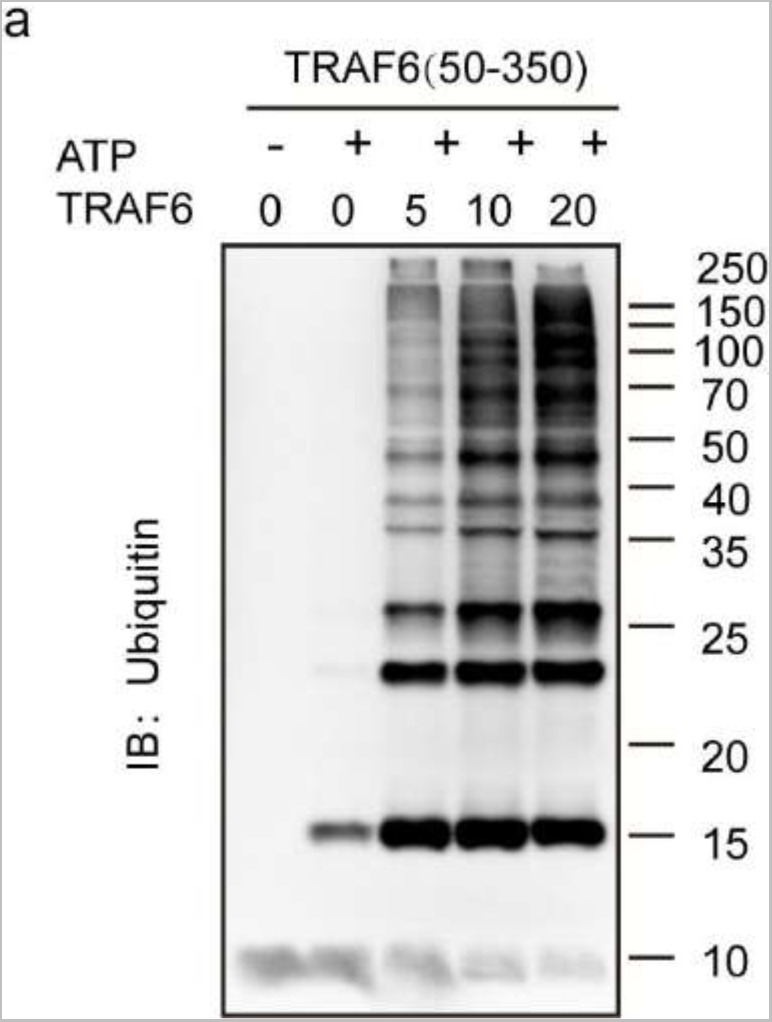
TRAF6 promotes the poly-ubiquitin chain synthesis in a concentration-dependent manner. A. Western blotting of TRAF6 ubiquitination reactions in vitro.

**Figure EV4.**
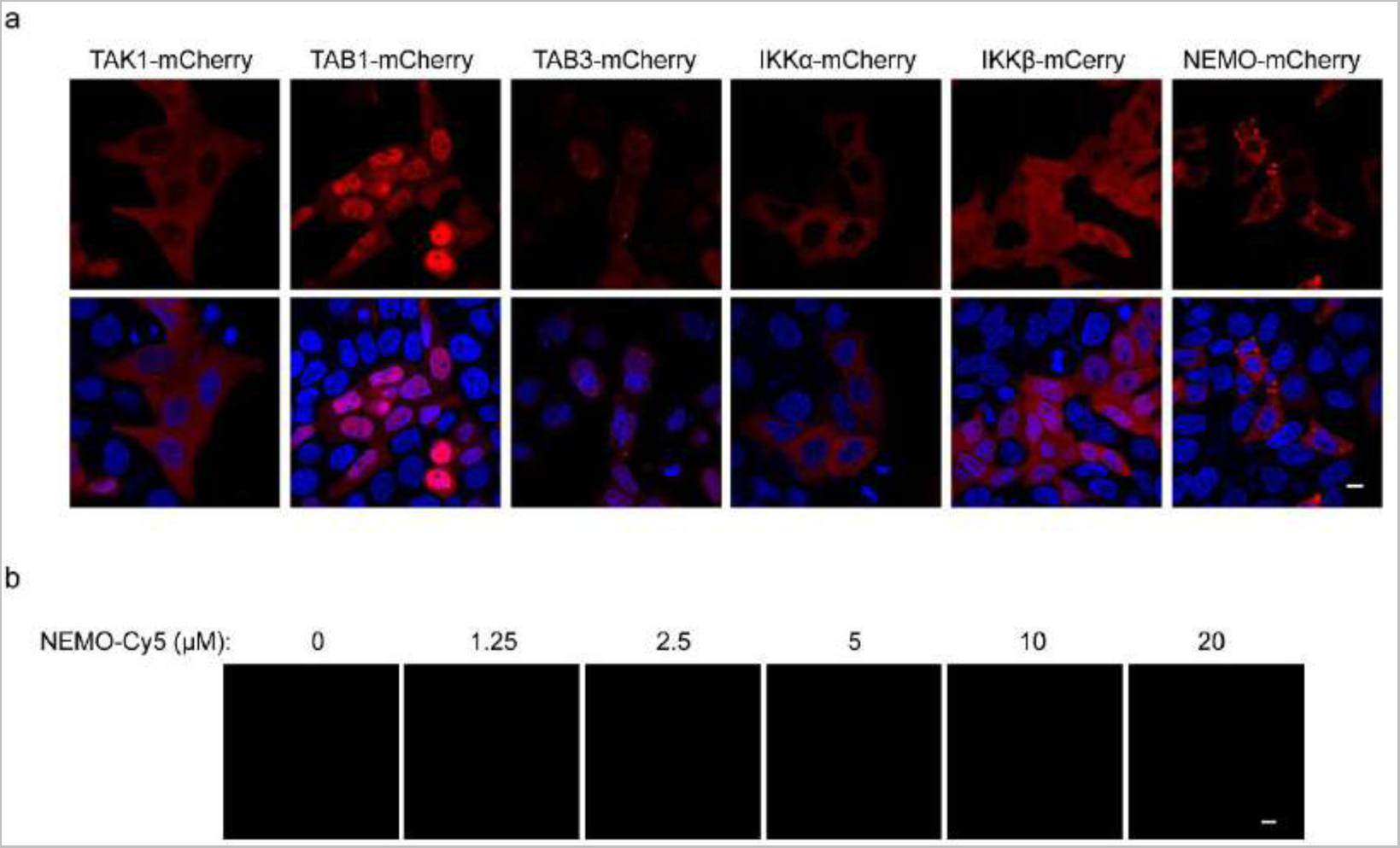
Downstream proteins can’t undergo phase separation without TRAF6. A. Representative images of TAK1-mCherry, TAB1-mCherry, TAB3-mCherry, IKKα-mCherry, IKKβ-mCherry and NEMO-mCherry overexpressed in HeLa cells alone.
B. Representative images of NEMO-Cy5 in phase separation buffer at indicated concentrations. Data information: Scale bars, 10 μm in (A) and (B).

**Figure EV5.**
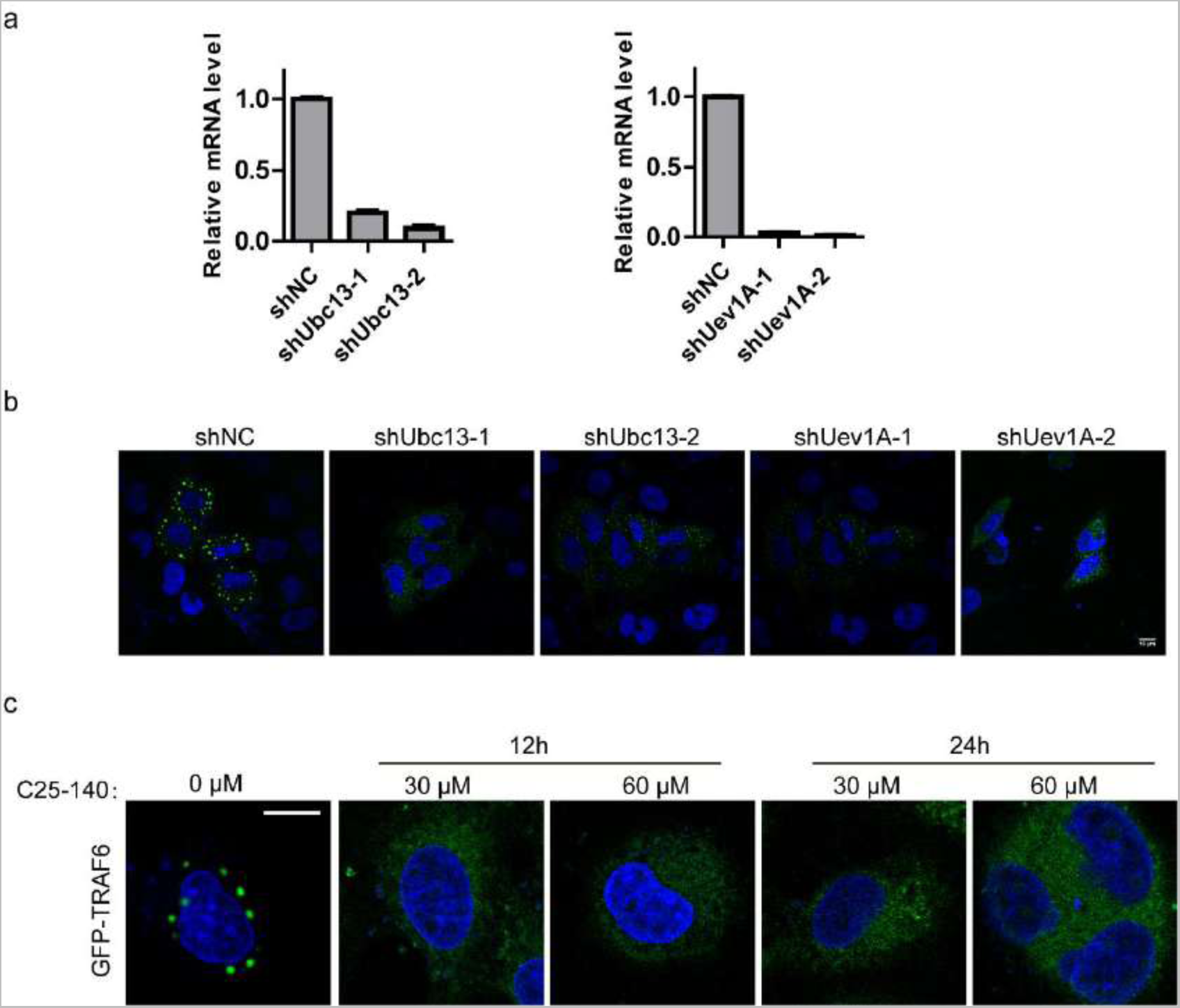
The E2 pair UBc13/Uev1A is important to the TRAF6 phase separation. A. ShRNA knockdown efficiency of Ubc13, Uev1A in HeLa cells.
B. Images of EGFP-TRAF6 overexpressed in shNC, shUbc13-1, shUbc13-2, shUev1A-1, shUev1a-2 HeLa cells.
C. Images of EGFP-TRAF6 in cells treated with different C5-124 concentrations for the indicated times. Data information: Scale bar, 10 μm in (B) and (C).

**Figure EV6.**
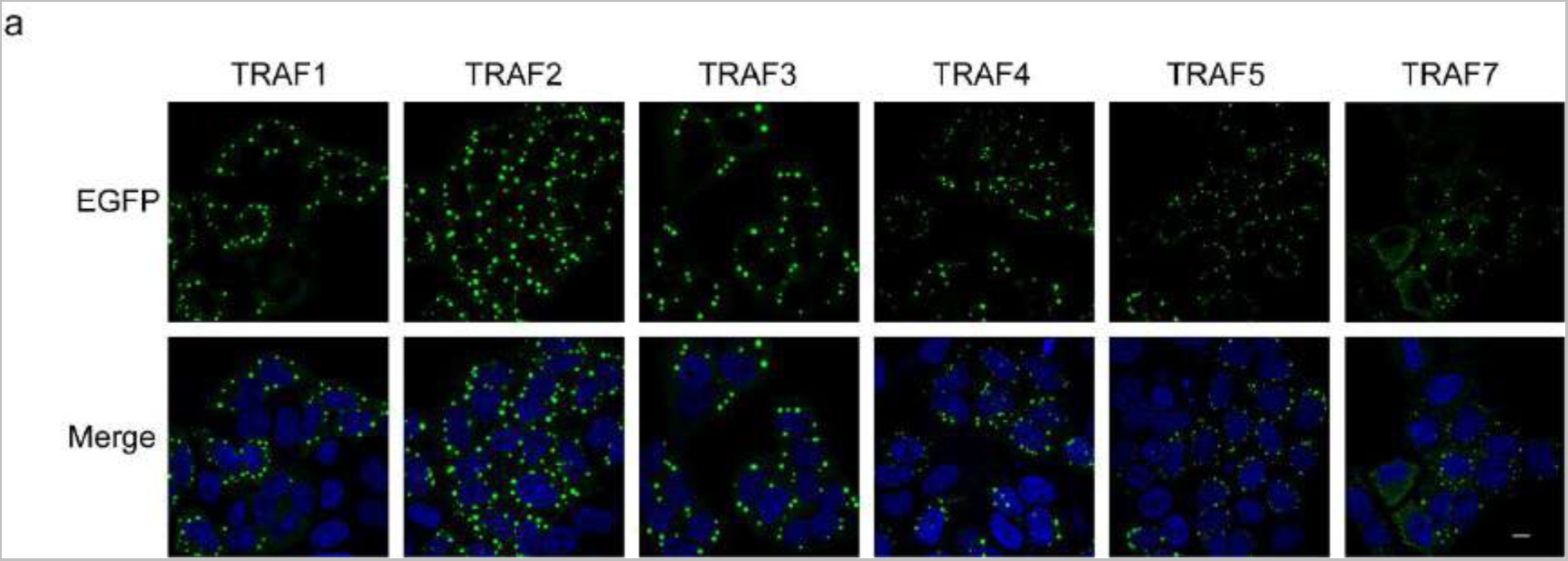
TRAF family proteins form droplets in cells. A. Images of EGFP-TRAF1, EGFP-TRAF2, EGFP-TRAF3, EGFP-TRAF4, EGFP-TRAF5 and EGFP-TRAF6 overexpressed in HeLa cells.

